# Applying integrated population models to guide conservation planning: Western Capercaillie in Scotland

**DOI:** 10.64898/2026.01.13.695102

**Authors:** Holly I. Niven, Fergus Blyth, Jack A. Bamber, Molly Doubleday, Steven R. Ewing, Kathy Fletcher, Sue Haysom, Kenny Kortland, Xavier Lambin, Robert Moss, Chris Sutherland, Laura J. Zantis, Jason Matthiopoulos

**Affiliations:** School of Biodiversity, One Health and Veterinary Medicine, University of Glasgow, Glasgow G12 8QQ, Scotland, UK; Independent Researcher; School of Biological Sciences, University of Aberdeen, AB24 2TZ Aberdeen, Scotland, UK; RSPB Centre for Conservation Science, Edinburgh, UK; GWCT, Hopetoun Estate Office, South Queensferry, EH30 9SL; NatureScot, Great Glen House, Leachkin Road, Inverness, IV3 8NW; Cairngorms Connect, Aviemore, UK; Forestry and Land Scotland, Smithton, Inverness, UK; Station House, Crathes, Banchory, Aberdeenshire; Centre for Research into Ecological and Environmental Modelling, University of St Andrews, St Andrews, UK; Institute of Environmental Sciences, Leiden University, P.O. Box 9518, 2300 RA Leiden, The Netherlands

**Author notes:** Deceased (Sue Haysom).

**Keywords:** Breeding success, climate-demography relationships, demographic modelling, ground-nesting birds, imperfect detection, lek attendance, population dynamics, vole cycles

## Abstract

1. Many conservation interventions prove ineffective because they lack a rigorous evidence-base. Multifactorial drivers and data limitations often hinder anticipatory planning and are difficult to tackle with standard methods. Predictive modelling approaches that integrate diverse data sources with biological hypotheses can bridge this gap by clarifying the drivers of decline and evaluating management options under uncertainty.
2. We developed a Bayesian integrated population model (IPM) applicable to lekking species, combining incomplete data from different life stages and seasons while accounting for observation error. IPMs are widely recognised for reducing bias relative to single-data-stream analyses, yet their application in conservation planning remains limited.
3. We applied the model to the threatened Western Capercaillie population in Scotland, which has been declining since at least the 1980s despite decades of conservation efforts, a typical case requiring urgent, evidence-based management. By fitting the model to 30 years of biased and incomplete monitoring data, we investigated associations between demographic processes and their potential drivers, including weather variables, fence collision mortality and a proxy of predation pressure. We used model predictions to evaluate the joint effectiveness of proposed management actions aimed at improving vital rates, including fence management and diversionary feeding.
4. Data integration improved population estimate precision by 17-49% relative to standalone national survey estimates, confirming that the decline continued from 1990-2023. Reproductive rate was related to the pattern of April warming, negatively affected by pre-breeding precipitation and positively by vole abundance, the latter consistent with the alternative prey hypothesis. Model predictions indicated that combining diversionary feeding with fence removal produced the most favourable conservation outcomes, but growth remained limited and uncertainty included possible continued decline.
5. 5. This modelling approach forms a key component of Scotland’s Capercaillie Emergency Plan 2025-2030, guiding management by assessing population-level responses and predicting conservation intervention outcomes. In future, the model could support formal adaptive management through iterative evaluation of interventions as new data become available. Beyond this case study, our integrated approach offers a transferable framework for managing multifactorial declines in other threatened species under uncertainty.

## 1 Introduction

Designing effective, evidence-based interventions for declining species is often constrained by sparse demographic and covariate data, cryptic life stages, multiple and indirect environmental drivers, and ecosystem stochasticity (Martin et al., 2022; McDonald-Madden et al., 2010; Schmolke et al., 2010). These challenges can lead to ineffective actions and wasted resources (Binley et al., 2025). Overcoming these limitations requires optimal inference via the formal integration of diverse data types to test biological hypotheses and evaluate explicit management actions under uncertainty (Bolam et al., 2023; Bradbury et al., 2001).

Integrated population models (IPMs; Schaub & Abadi, 2011), combining a biological process model with an observation model, enable the integration of population survey and demographic data while accounting for uncertainty, improving parameter estimation and reducing bias compared to single-data-stream approaches (Frost et al., 2023; Schaub & Abadi, 2011). However, their application in conservation remains limited, due to their custom nature and technical complexity (Frost et al., 2023). IPMs are particularly valuable when key demographic processes are only partially observed, as in lekking species where monitoring primarily targets displaying males (Walsh et al., 2010).

The Western Capercaillie (*Tetrao urogallus*; henceforth capercaillie) illustrates why integrated approaches are invaluable for conserving species with complex life histories. Populations have declined across much of its range south of the boreal zone in Europe, including Cantabria (Jiménez et al., 2022), the central Pyrenees (Gil et al., 2020), and the Black Forest in Germany (Coppes et al., 2016). In Scotland, the species is red-listed (Stanbury et al., 2021) and has disappeared from much of its historical range. Despite decades of conservation efforts, estimated numbers have declined from ≈20,000 in the 1970s (Moss et al., 2024) to 2,200 (95% CI: 1,500-3,200) in 1992-1994, and 532 (95% CI: 227-810) in 2021-2022 (Catt et al., 1998;

Wilkinson et al., 2023). As well as periodic national surveys, monitoring in Scotland comprises annual brood counts (chicks and hens) and spring lek counts. Like most ecological datasets, these have historically been analysed separately (Baines et al., 2016; Moss et al., 2000, 2001) or only partially integrated, leading to discrepancies with estimates from population counts (Baines & Aebischer, 2023) and underscoring the need for a holistic modelling approach to support management decisions.

As with many threatened species, the decline of Scottish capercaillie involves multiple indirect and cumulative drivers that are likely to confound univariate analyses (He & Callaway, 2009). Low breeding success was likely the primary demographic cause of the decline. In particular, climatic changes are thought to have played a role (Baines et al., 2011; Moss et al., 2001), by reducing hen condition (Coppes et al., 2021; Slagsvold & Grasaas, 1979) and chick survival (Baines et al., 2016; Picozzi et al., 1999; Wegge et al., 2022). The decline may have also been exacerbated by mortality from collisions with fences (Catt et al., 1994) which persists due to newly erected, remaining or poorly visible fences. Capercaillie are also vulnerable to egg depredation by changing mesocarnivore communities including recovering pine marten (*Martes martes*) (Baines et al., 2016; Bamber et al., 2024; Palencia & Barroso, 2024) and other predators such as red fox (*Vulpes vulpes*), and European badger (*Meles meles*). Predation pressure may intensify in years when keystone prey such as field voles (*Microtus agrestis*) (Caryl et al., 2012; Leckie et al., 1998), decline (an indirect effect known as the alternative prey hypothesis; Angelstam et al., 1984; Kjellander & Nordström, 2003). Changes in grassland vole population cycles, such as the dampening of cycle amplitude observed across Europe over the past two decades (Cornulier et al., 2013) may have altered predation dynamics, potentially reducing breeding success.

Scotland’s Capercaillie Emergency Plan 2025-2030 outlines measures to improve breeding success and survival across the species’ remaining core range within the Cairngorms National Park (NatureScot & Cairngorms National Park Authority, 2024). The plan advocates ongoing actions such as fence marking and removal, lethal predator control, and habitat management (Kortland & Capercaillie BAP Group, 2006; Moss & Picozzi, 1994; Watson & Moss, 2008; Wilkinson et al., 2023), which have not led to recovery (Wilkinson et al., 2023). The plan also discusses novel promising approaches (Bamber et al., 2024); diversionary feeding deploys supplementary food sources to divert mesopredators from capercaillie during the breeding season (Kubasiewicz et al., 2016). Although experimental application can increase breeding success by ≈130% (Bamber et al., 2025b), the efficacy of these measures for mitigating population declines is unknown.

To improve understanding of drivers of decline and inform management, we developed a Bayesian state-space IPM that combines multiple data sources to estimate trends in capercaillie abundance and vital rates. We use this model to quantify the direction and magnitude of hypothesised drivers of decline and to predict how multiple management actions may influence future population viability, providing evidence to guide conservation planning. Our approach is applicable to many species facing multifactorial declines to enhance insights from biased, incomplete, or fragmented datasets.

## 2 Materials and Methods

### 2.1 Data

#### 2.1.1 Study area and demographic data

Demographic data were from the Badenoch & Strathspey (B&S) region (Figure 1), the species’ main stronghold in Scotland containing ≈80% of the population in 2021 (Wilkinson et al., 2023). We used three sources of population data:

1. Spring counts of males (*l*_i,t_), in lek *i* and year *t*. A total 67 leks were surveyed over 27 years between 1990 and 2020 (Baines & Aebischer, 2023; Picozzi et al., 1992), with a median of 28 leks per year (range 1-60) (Fig. S1). Lek counts likely miss some younger males and absentee adults (Watson & Moss, 2008) (see Section 2.2.1 for age definitions). Each lek was assigned to one of 25 forest areas (Fig. 1), denoted *f* (hereafter forest), when assigning covariate values at the forest area scale (see Section 2.1.2), with median 1 (range 1-13) leks per forest area.
2. Late summer brood counts of hens (ℎ_*f,t*_) and chicks (*c*_*f,t*_), were collected using pointing dogs. 19 of 25 forests were surveyed over 30 years between 1990 and 2020 (Baines et al., 2004; Baines & Aebischer, 2023), with a median of 7 forests per year (range 1-14) (Fig. S2). Observed reproductive rate 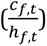 showed no linear temporal trend (Supplement, SI 1).
3. National survey counts provided population estimates (*P*_st_) for the 2009-2010, 2015-2016 and 2020-2021 winters (Table 1) that accounted for observation error (Ewing et al., 2012; Wilkinson et al., 2018, 2023). Sex-specific estimates were not available for separate regions.

**Figure 1:**
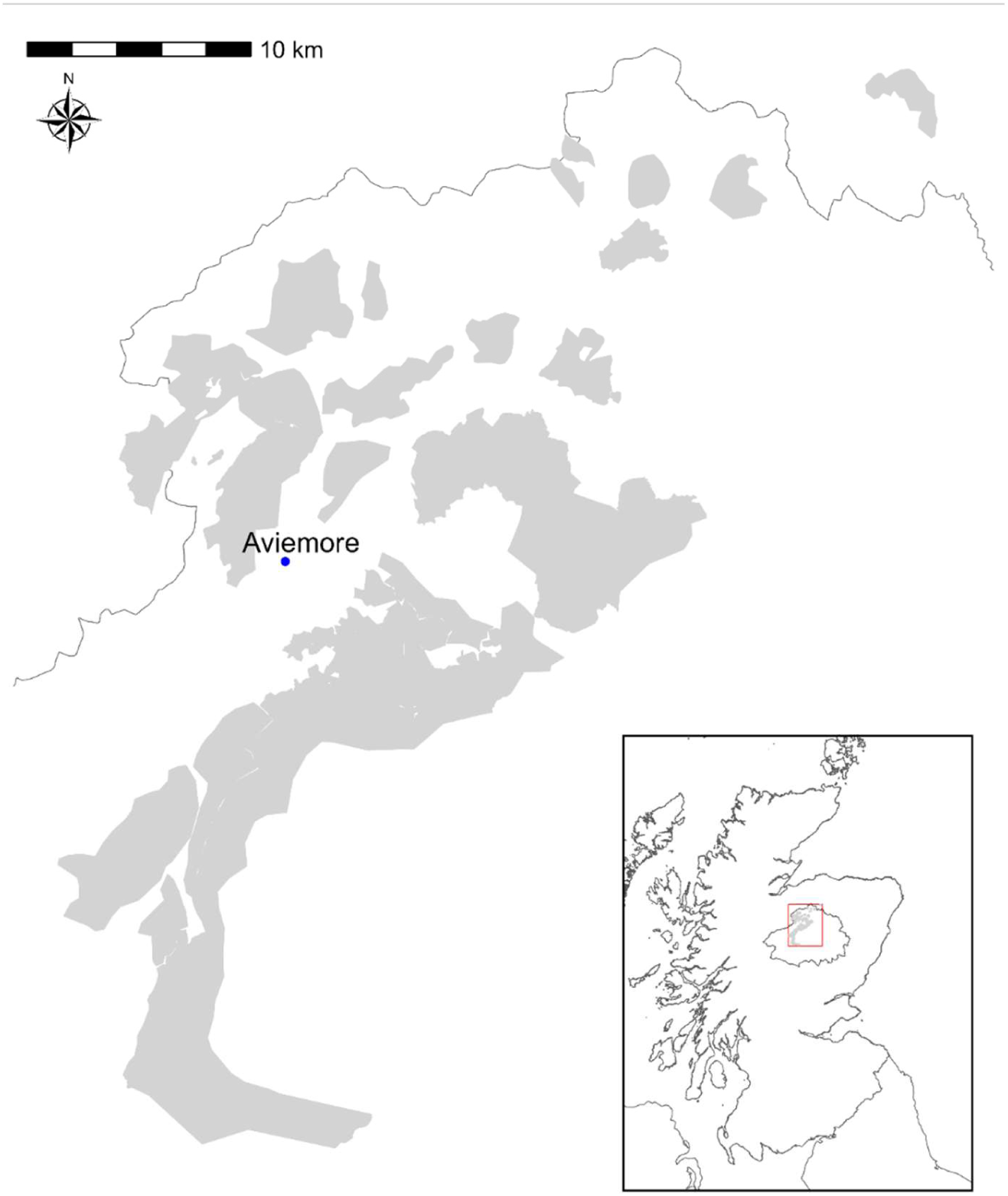
Map of the study area showing approximate forest areas (grey) with capercaillie presence between 1990 and 2023. The thin black line indicates the Cairngorms National Park boundary within Scotland (inset). The blue dot marks Aviemore, used as the location for weather data (Section 2.1.2).

**Table 1:** National survey counts from Badenoch and Strathspey (B&S) with the winter conducted, median and lower and upper confidence limits (CLs), and proportion of males. For 2015-16 and 2021-2022, the proportion of males is based on sex-specific estimates for the entire capercaillie range (none were available for B&S), whereas for 2009-10 it relies on observed data for the whole range, as there were no sex-specific population estimates.

| Year | Median | Lower CL | Upper CL | Proportion of<br>males |
| --- | --- | --- | --- | --- |
| 2009-2010 | 968 | 578 | 1514 | 0.45 |
| 2015-2016 | 976 | 711 | 1319 | 0.46 |
| 2021-2022 | 423 | 202 | 586 | 0.54 |

#### 2.1.2 Covariate Data

Candidate covariates chosen for data availability and biological relevance (Table 2) included:

1. Fence density (km^−1^) (γ_i_) in 2020: total length of fences within 1km of the centroid of all recorded locations for each lek. Interannual fence data were unavailable, so densities were assumed constant at 2020 levels.
2. Weather covariates: pre-breeding precipitation, mean April temperature, an index of April warming and June precipitation days for 1990-2023, derived from Global Surface Summaries of the Day (GSOD) data for Aviemore weather station (NOAA National Centers of Environmental Information, 1999) using the GSODR package (Sparks et al., 2017) (Fig. S3, S4, S5, S7). Spring snow cover indices 1990-2022 (Spracklen & Spracklen, 2023; Fig. S6) were used instead of unavailable snow melt data. The 2023 snow cover value was predicted (SI 2).
3. Pine marten scats (Scats km^-1^ day^-1^) (*P*_m f,t_) in May-June: collected on tracks in 5-7 forests in 1995, 2009 and 2013 (Baines et al., 2013; Summers et al., 2015).
4. Spring vole abundance indices (*V*_*f,t*_): derived from annual quadrat surveys in five forests (2018-2023), using the proportion of quadrats with field vole (*Microtus agrestis*) signs (Lambin et al., 2000). Field voles are the primary preferred prey of red foxes and pine martens in Scotland (Caryl et al., 2012; Leckie et al., 1998), and thus serve as proxies of preferred prey availability for mammalian mesocarnivores. To ensure plausible estimates for missing B&S vole values, we used long-term vole sign indices from Kielder Forest (1990–2023; Lambin et al., 2000) as a reference series during reconstruction (see Section 2.2.4).

**Table 2:** Demographic and covariate data used in the analysis, including (where applicable): data name, temporal coverage, spatial scale, calculation method, unit, and hypothesised effect (SI 4). *T*_3_, *T*_2_ and *T*_1_ refer to the mean temperature during the last, middle and first 10 days of April, respectively, based on observed daily means. RR denotes reproductive rate. Covariate hypotheses are classified according to Wegge et al., 2022 taxonomy.

| Demographic and covariate data | Temporal Coverage | Spatial scale applied at | Unit | Calculation | Hypothesis |
| --- | --- | --- | --- | --- | --- |
| National survey | 2009-10,<br>2015-16,<br>2021-2022 | B&S | Count (no unit) | NA | NA |
| Lek data | 1990-2020<br>(Fig. S1) | Lek (each lek located within a forest) | Count (no unit) | NA | NA |
| Brood data (counts of hens and chicks) | 1990-2020<br>(Fig. S2) | Forest | Count (no unit) | NA | NA |
| Pre-breeding precipitation | 1990-2023 | B&S | mm | Sum of daily precipitation in March | Hen condition hypothesis: More precipitation has a -ve effect on RR (Coppes et al., 2021) . |
| April temperature | 1990-2023 | B&S | °C | Mean of observed mean daily temperatures | Chick food hypothesis: Higher April temperature has a -ve effect on RR (Baines et al., 2016). |
| April warming index | 1990-2023 | B&S | °C | $\frac{[(T_2 - T_1) - (T_3 - T_2)]}{2}$ (Moss et al., 2001) | Hen condition hypothesis: Larger temperature rise in early April has a +ve effect on RR (Moss et al., 2001). |
| June precipitation days | 1990-2023 | B&S | Count (no unit) | No. precipitation days in days 1-20 | Chick weather hypothesis: More precipitation days has a -ve effect on RR (Picozzi et al., 1999; Summers et al., 2010) |
| Snow cover | 1990-2022 | B&S | No unit | Difference in mean snow cover May to mid-September (of 9 high-altitude sites across the Scottish Highlands calculated from remote sensing data) from normal over study period (Spracklen & Spracklen, 2023) | Hen condition hypothesis: Snow cover could have a +ve or -ve effect on RR (Slagsvold & Grasaas, 1979; Wegge & Rolstad, 2011) . |
| Voles | 2018-2023 | Forest | Proportion (no unit) | See section 2.1.2 | Predation hypothesis: More voles have a +ve effect on RR (Kjellander & Nordström, 2003; Wegge et al., 2022) . |
| Pine Martens | 1995, 2009,<br>2013 | Forest | Scats/km/d<br>ay | See section 2.1.2 | Predation hypothesis: More pine martens have a -ve effect on RR (Baines et al., 2016; Bamber et al., 2024) . |
| Fences | 2020 | Forest | km | See section 2.1.2 | More fences have a -ve effect on first-year female survival (Moss et al., 2000) . |

Of those covariates, only April warming index (SI 1; Table S1) displayed an increasing linear trend, contrary to the historical decreasing trend reported by Moss et al. (2001) between 1975-1998. Variance inflation factors (all < 2) indicated no problematic collinearity among predictors, including with time. Covariates were z-standardised to aid model convergence and comparability of effect sizes for each demographic rate (Schielzeth, 2010). Pine marten indices were kept unstandardised to allow modelling of density-dependent population growth (see SI 5).

### 2.2 Model Description

The model was fitted for 1990-2023, to incorporate all demographic and covariate data. IPMs comprise two components: a process model describing biological processes assumed to govern the system’s true (“latent”) states, and an observation model linking these latent states to data while accounting for observation error (Auger-Méthé et al., 2021) (Fig. 2). Full details of priors, model symbols, running specifications, and covariate selection are provided in Supplementary Information (SI 4; Table S2; SI 6).

**Figure 2:**
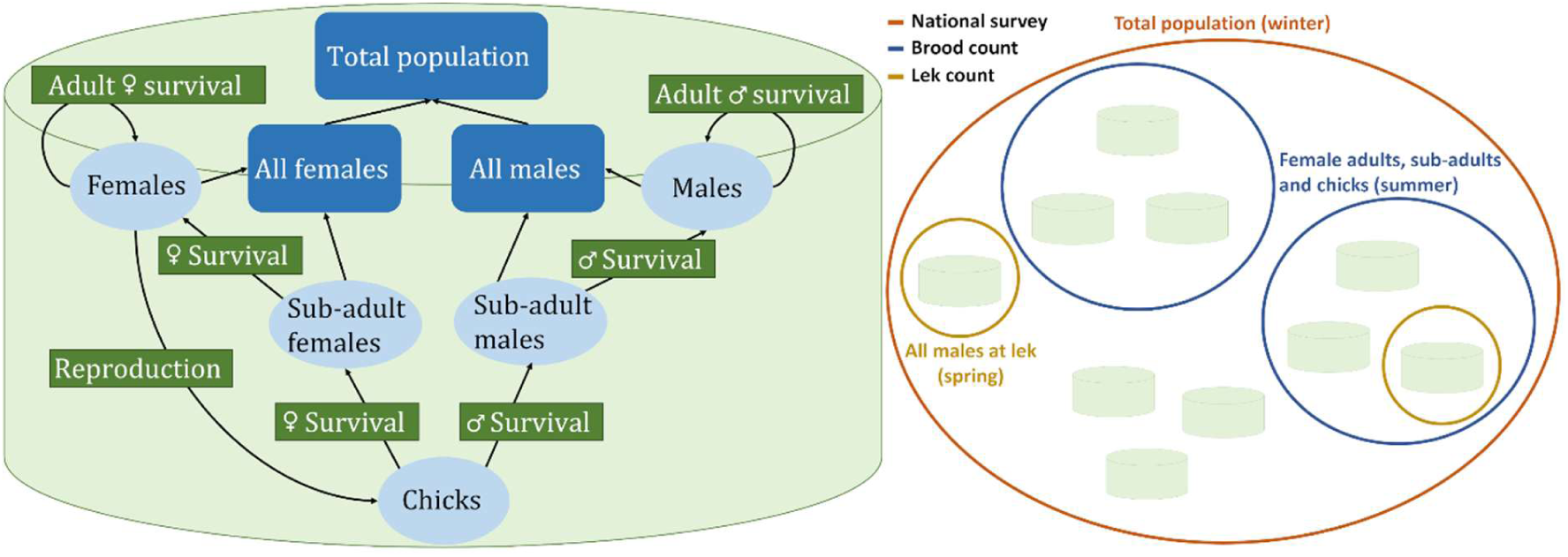
Structure of the Bayesian state-space integrated population model for capercaillie (without covariates). Left: Simplified diagram of the sex- and age-structured process model, showing population states (light blue ovals), demographic processes (survival and reproduction; green rectangles) and monitored variables (dark blue rounded rectangles). Right: Stylised representation of the observation model, illustrating the different spatial scales, spatial imbalance and seasonal timing of monitoring data incorporated into the model. Data streams include national surveys (whole population, winter; orange circle), brood counts (adult and 14-month females and chicks in some forests, summer; blue circles), and lek counts (males at monitored leks, spring; yellow circles). Cylinders represent lek subpopulations. See Sections 2.2.2 and 2.2.3 for details of process and observation models.

#### 2.2.1 Definitions and assumptions

We modelled each lek separately to capture inter-lek variation in demographic rates. Each lek was assumed to have uncounted individuals (males not at the lek and all classes of females) that were imputed by the model (see Section 2.2.3; Fig. 2). Forest-specific covariates and brood counts were assigned to each lek in a forest. Given the declining population, new lek formation after 2020 was considered unlikely. Leks can shift over time, so a relocated lek may occasionally be recorded as new; however, this is unlikely to affect our model because previous locations appear as zero in the data. Because the population is declining, we assumed no migration between leks, treating any dispersal exchanges as symmetric, on average. However, immigration could reduce extinction risk for small leks (rescue effect; Brown & Kodric-Brown, 1977), so we explored sensitivity to potential rescue effects in our population projections (see Section 2.3; 3.3).

Late April/early May lek counts form the temporal reference for the model. Observations at other times, national surveys (January midpoint) and brood counts (August), were linked to latent states by applying additional mortality between May and the observation month.

The sex ratio (*p*_C_) was estimated as a constant with a female bias (Moss et al., 2000). Adult females were defined as >23 months old, assuming hatching in early June, and maturity after lek counts. Brood counts include 14-month hens, adult hens (>26 months old), and chicks. 14-month hens are less likely than adults to successfully rear chicks, especially when overall breeding success is poor (Fletcher & Baines, 2020). We therefore estimate reproductive rate as chicks per adult hen using latent states in the model, while also reporting chicks per hen (including 14-month hens) for comparison with published estimates. Males were divided into three subadult and one adult stages (Fig. 2). First-year individuals experience first-year (juvenile) survival and older stages share adult survival.

#### 2.2.2 Process Model

We modelled population dynamics at each lek using age- and sex-structured classes linked by survival and reproduction. Each lek *i* in year *t* contained adult females (𝐹_i,t_), adult males (𝑀_i,t_), subadult female classes (𝑓_a i,t_, a ∈ {0,1}) and male classes (𝑚_a i,t_, a ∈ {0,1,2})(Fig. 2). (Full notation in Table S2).

For comparison with national surveys, total population size in winter (January midpoint; 8 months after May lek counts) and the proportion of males in the population were calculated as:

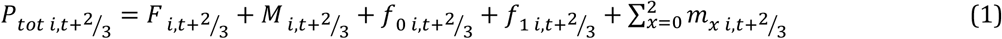

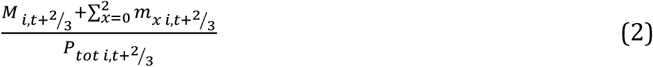

and summed over all leks.

This formulation accounts for mortality between spring and winter (SI 3).

Age transitions were modelled as Poisson processes based on chick production and stage-specific survival:

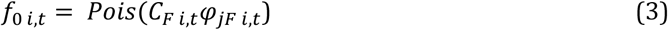

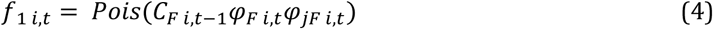

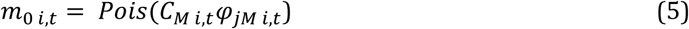

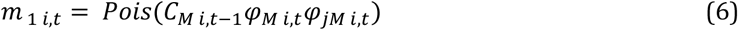

where 𝐶_x,i,t_ is the number of chicks of sex 𝑥 ∈ {𝐹, 𝑀}, *ф*_jx,i,t_ is first-year survival, and *ф*_x,i,t_is adult survival. Second-year males were modeled similarly:

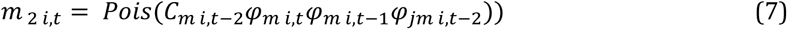

Adult numbers were the combination of survivors (𝑆_x i,t_) and new recruits:

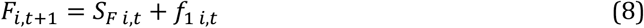

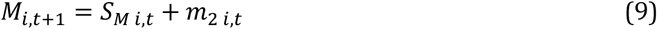

Survival was modelled as a binomial process:

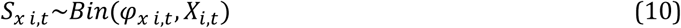

with sex specific survival probabilities *φ*_x i,t_.

First-year (juvenile) and adult survival probabilities were modelled using logistic functions with lek and year random effects and a fence density covariate (γ_i_):

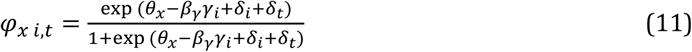

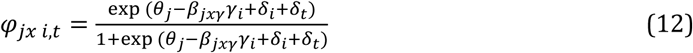

Adult survival had sex-specific intercepts (𝜃_x_) (Augustine et al., 2020) and a shared fence mortality coefficient (𝛽_γ_). First-year survival (φ_jx i,t_) shared an intercept (𝜃_j_), but included sex-specific fence mortality coefficients (𝛽_jxγ_), reflecting that first-year females may be particularly impacted during natal dispersal (Moss et al., 2000). Random effects for lek and year (𝛿_i_, 𝛿_t_∼𝑁(0, 𝜎^2^)) account for unmodelled variation among leks and between years that is not captured by the covariates or other process model components.

Reproductive rate (𝜇_i,t_, chicks per adult hen) was modelled as:

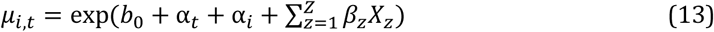

where 𝑏_0_ is the baseline reproductive rate, αi, αt ∼𝑁(0, 𝜎^2^) are random effects per lek and year (also see Eq. 11,12), and *Χz*, are covariates (Section 2.1.2; Table 2), with coefficients 𝛽_z_.

Total chicks produced, 𝐶_i,t_, followed a Poisson distribution based on the expected number of adult females in August (3 months after May lek counts), 𝐹_i,t+_1_/_ reproducing at per-capita rate,

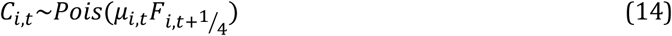

Extra-Poisson dispersion is accounted for by the random effects. Chicks-per-hen ratio was calculated as:

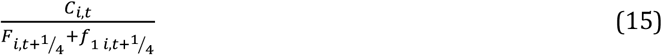

Chicks were split by sex using a binomial draw with sex-ratio *p*_C_:

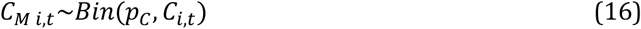

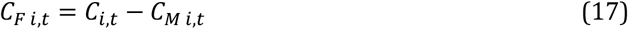

#### 2.2.3 Observation model

We constructed observation models for each data source to account for imperfect detection, seasonal mortality between observations, and observed life stages (Fig. 2).

We modelled the observed National survey count (*P*_s t_) as a Gaussian process centred at the true (unobserved) population size across leks, scaled by detection probability (*p*_s_):

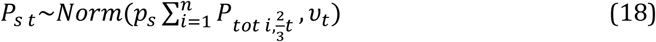

where u_t_ is the standard error of the published national survey point estimates. The Gaussian likelihood reflects the central-limit tendency of the aggregated national survey counts.

Observed lek counts (*li,t*) were modelled as a Poisson process, appropriate for discrete count data, allowing for the proportion of lek attendance (*p*_l_):

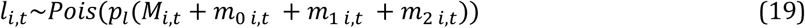

Brood counts were spot checks, so they could only inform breeding success. Observed chick counts were modelled conditional on the number of hens observed in each forest (ℎ_*f,t*_), adjusted for the proportion of adults and the forest mean reproductive rate. Because hen counts may include both adult hens and 14-month hens, we first derived the proportion of adult hens in August (*p*_h f,t_), to estimate chicks/adult hen, by averaging across all leks (𝑦_f_) within each forest:

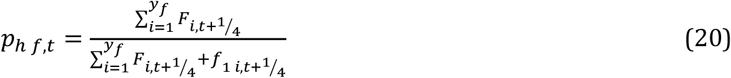

Forest area reproductive rate (𝜇_*f,t*_) was calculated as the mean across leks to match the spatial scale of brood counts:

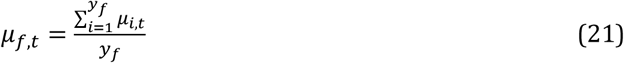

Observed chick counts (*c*_*f,t*_) were then modelled as a Poisson variate:

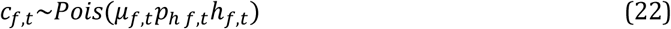

#### 2.2.4 Modelling predator effects

Vole cycles in Kielder were synchronous with those in B&S during overlapping years (Fig. S8) and consistent with European-scale dynamics (Cornulier et al., 2013). Observed B&S reproductive rate cycles showed a 3–4 year cycle, largely corresponding to Kielder vole cycles (Fig. S9). We therefore modelled standardised vole indices for each B&S forest (*V*_*f,t*_) as a normal distribution centred on the standardised Kielder vole index for that year (*V*_K,t_):

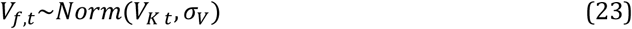

where 𝜎_V_ is the standard deviation representing the expected variability in local vole indices around the Kielder vole index for a given year (SI 4; Table S2).

Pine martens were eventually excluded from model selection because their limited data availability impaired model convergence. See SI 5 for full details of modelling approach and SI 6 for model selection procedure.

### 2.3 Simulated projection

A forward simulation without covariates was implemented in R using 1,000 samples from an MCMC chain thinned by 10, initialised from the last two years of model output and projected to 2070. Two scenarios were run: (1) baseline without rescue and (2) rescue-effect, where leks losing all adult males or females could gain one individual with probability 1/67 (inverse of total leks). For each scenario, adult female survival (−30% to +30%) and reproductive rate (−50% to +450%) were varied relative to posterior medians (logit^−1^(𝜃_F_) = 0.70; exp (𝑏_0_) = 0.70 chicks per adult hen), with changes defined on the probability scale for survival and original scale for reproductive rate, then transformed to link scales (logit, log) for implementation. The same adjustments were applied to male and first-year survival using their respective baselines. These ranges reflect plausible demographic responses, including those achievable via management.

To prevent unrealistic growth, we capped the total number of males or females per lek at 50 (Watson & Moss, 2008), scaled by attendance (*p*_l_). For each scenario, we calculated (1) probability-of-no-return in 2060 (percentage of simulations where the female population<75, range: 50–100; Moss, 2001), and (2) mean annual population growth over 2023–2033. The no-return threshold is assumed to represent a point beyond which population recovery is unlikely due to demographic and genetic effects of small population size (Moss, 2001). For comparison with Baines & Aebischer (2023), we also report extinction probability (≤ 1 male) in 2045 and 2070 for the *status quo* (i.e. no change) scenario.

Finally, empirical intervention effects were overlaid onto the simulation outputs: survival gains from fence marking (6.70%, 95% CI: 1.37-15.3%; Baines & Andrew, 2003) and removal (11.0%, 95% CI: 2.92-19.8%; Moss et al., 2000), and reproductive rate gains from diversionary feeding (130%, 95% CI: 44.1-371%; Bamber et al., 2025b), estimated via bootstrapping (SI 7).

## 3 Results

### 3.1 Model selection

Model selection identified voles (probability of inclusion, 𝐼_j_ = 1), April warming index (𝐼_j_ = 0.79) and pre-breeding precipitation (𝐼_j_ = 0.50) as the most influential covariates on capercaillie population dynamics (SI 6), with limited evidence for other covariates (Table 3). By magnitude, the median effect size of voles (positive) was approximately 1.5 times larger than that of April warming index (positive) and precipitation (negative) (Table 3).

**Table 3:** Model selection results showing the posterior mean probability of inclusion of each covariate (mean of the indicator variable 𝐼_j_), along with posterior median estimated effect sizes (𝛽_j_) and associated 95% credible intervals (CRIs) (SI 6). Covariates are ordered by 𝐼_j_.

|  | p(inclusion) | Median | Lower CRI | Upper CRI |
| --- | --- | --- | --- | --- |
| Voles | 1.000 | 0.463 | 0.367 | 0.560 |
| April warming index | 0.791 | 0.299 | 4.686E-05 | 1.629 |
| Pre-copulation precipitation | 0.505 | -0.288 | -7.469E-06 | -2.240 |
| Snow cover | 0.358 | -6.972E-03 | -0.148 | 0.111 |
| Fences (first-year male survival) | 0.345 | -0.586 | -1.492E-05 | -2.583 |
| June precipitation days | 0.205 | -0.465 | -2.102E-05 | -2.770 |
| Fences (first-year female survival) | 0.062 | -0.629 | -3.864E-06 | -2.922 |
| Fences (adult survival) | 0.022 | -0.672 | -1.260E-05 | -2.964 |
| April temperature | 0.017 | -0.673 | -7.694E-06 | -2.967 |

### 3.2 Population dynamics

Estimated population size declined markedly over time: from approximately 2985 individuals (95% CRI 2162-4098) in 1995 to 1262 (95% CRI 1024-1573) in 2005. The population then fluctuated around 1200 until 2015, before declining further to 689 (95% CRI: 421-1089) in 2023 (Fig. 3A). The credible intervals for our model-based estimates were 49.4%, 16.9%, and 26.3% narrower than the confidence intervals from the national surveys (Table 1, Fig. 3A). In survey years, median population estimates were slightly higher than national survey point estimates, but their 95% intervals overlapped. Similarly, the estimated observation error for surveys was *p*_s_= 0.96 (95% CRI: 0.88-1.04; Table S2), indicating no deviation from 1.

**Figure 3.**
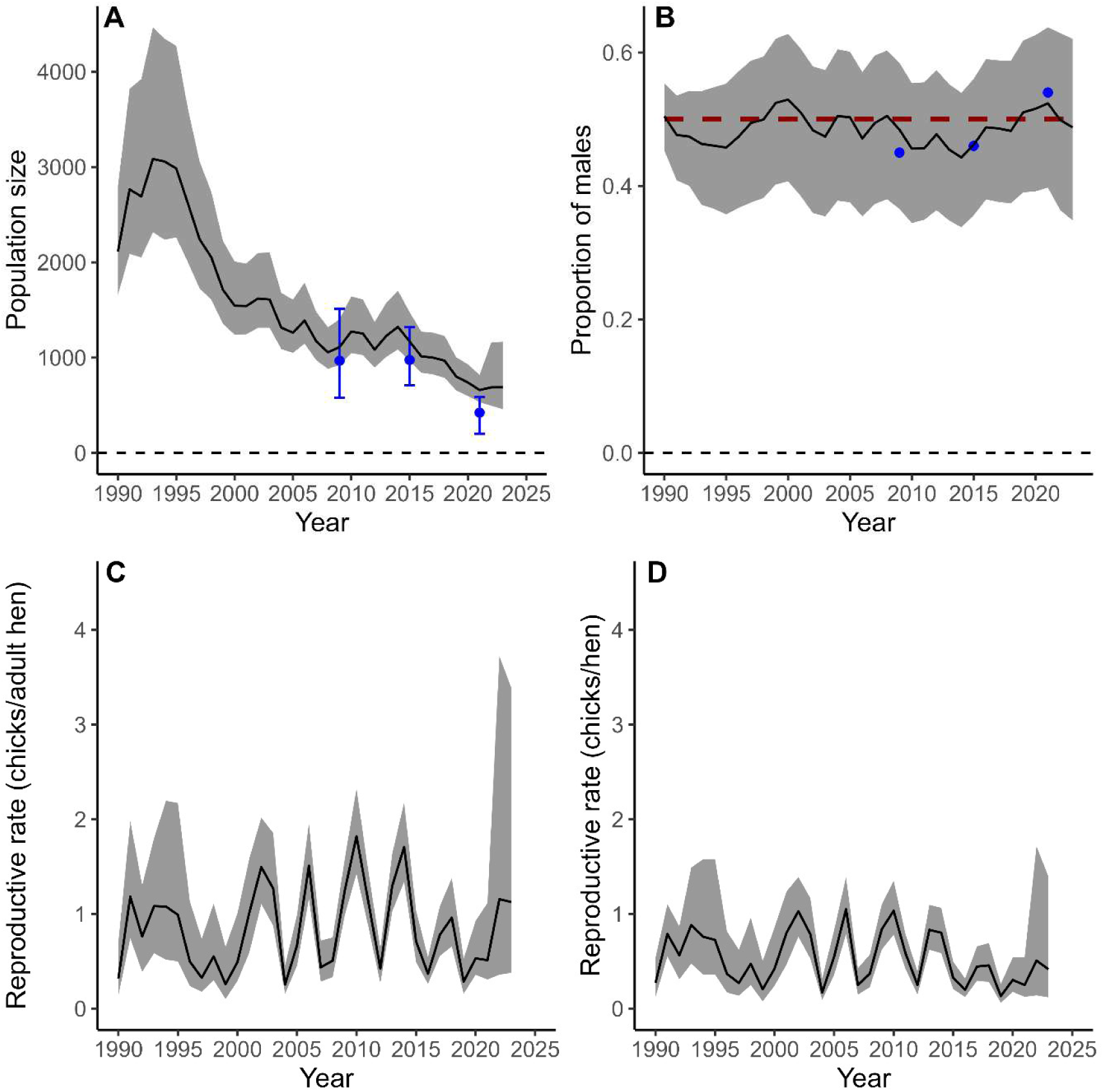
Modelled capercaillie population dynamics in Badenoch and Strathspey, Scotland from an IPM integrating national survey estimates, lek counts and brood counts. All panels show posterior medians (black lines) and 95% credible intervals (shaded areas). A: Estimated winter population size (all age classes). Blue points indicate national survey medians with survey-specific confidence intervals. B: Estimated proportion of males in winter; blue points are national survey point estimates for the whole Scottish range (Table 1), and the red dashed line indicates parity (0.5). C: Modelled reproductive rate expressed as chicks per adult hen (adult hens>23 months). D: Modelled reproductive rate expressed as chicks per hen (all hens).

The estimated proportion of all males seen at leks was *p*_l_= 0.34 (95% CRI: 0.28-0.40) (Table S2). The estimated proportion of males in winter closely matched national survey estimates (Fig. 3B). The estimated mean reproductive rates showed a 3-4 year cycle (Fig. 3C, 3D), coinciding with observed reproductive rate and largely with Kielder vole cycles (Fig. S10). There was no linear temporal trend in the estimated reproductive rates or temporal random effect (α_t_) (SI 1).

### 3.3 Projection with null model

Population projections for the period 2023-2033, under the *status quo,* forecast median annual decline (median= –3.46%, 95% CRI: -6.66%-4.92%) with a 31.2% (19.6-39.3%) probability-of-no-return in 2060 (Fig.4; Fig. 5). Using the Baines & Aebischer (2023) extinction criterion, our status-quo simulation estimated a 0% probability of extinction in 2045, and 1.6% in 2070.

**Figure 4:**
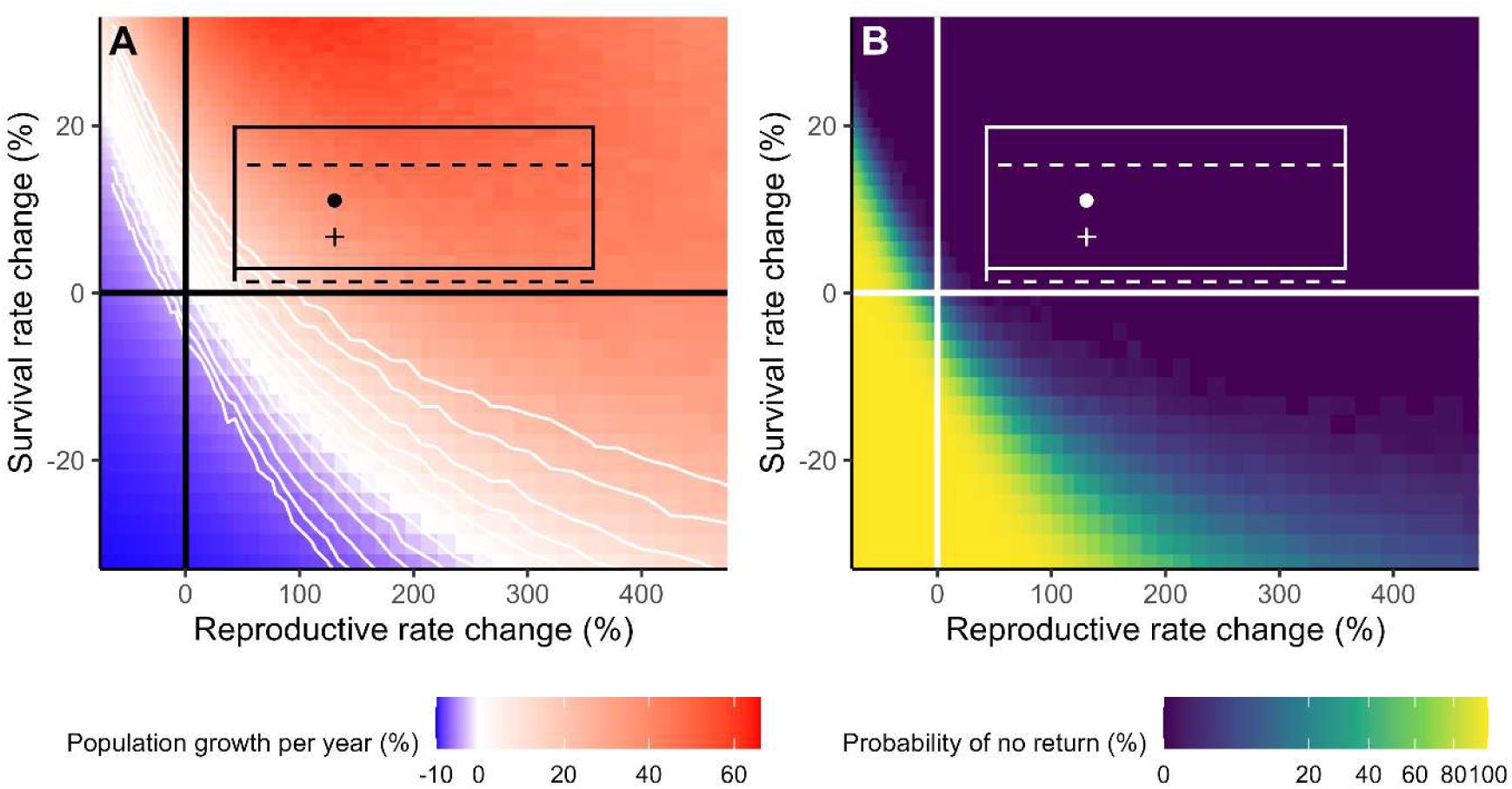
Predicted effects of changes in survival and reproductive rate on Scottish capercaillie population dynamics (2023-2060) based on the fitted population model. Outcomes of three management interventions are plotted in a two-dimensional space of changes in survival (y axis) and reproductive success (x-axis). Solid-line rectangles represent outcomes from fence removal and diversionary feeding; dots indicate median outcomes. Dashed lines indicate the range of outcomes for fence marking and diversionary feeding; crosses mark median outcomes. Changes are relative to the *status quo* simulation indicated by crosshairs (black in A, white in B) (median female survival = 0.70; reproductive rate chicks/adult hen, = 0.70). A: Median annual total population growth (2023-2033); white contours show quantiles: 5^th^-95^th^ in 10% increments. B: Probability-of-no-return in 2060 (population< 75 females).

Almost all simulations within management scenario outcomes (diversionary feeding increasing reproductive rate and fence marking or removal improving survival) fell within the “recovery zone” of the viability map (Fig. 4A), corresponding to near-zero extinction risk (Fig. 4B).

At the lower bounds of intervention effectiveness, diversionary feeding alone reduced extinction risk to almost zero (Fig. 5D) and was predicted to yield a slow and uncertain population growth (3.61%; Fig. 5B). Fence removal was more effective than marking: each still yielded negative median growth (Fig. 5A) but reduced the probability-of-no-return to 21.8% for marking and 12.5% for removal (Fig. 5C). Combining diversionary feeding with fence removal resulted in the highest median growth (7.47%; Fig. 5B). Growth rates for scenarios with and without rescue effects were very similar (Fig. 5A; Fig. 5B), while the median probability-of-no-return was slightly reduced when rescue effects were included (Fig. 5C).

**Figure 5:**
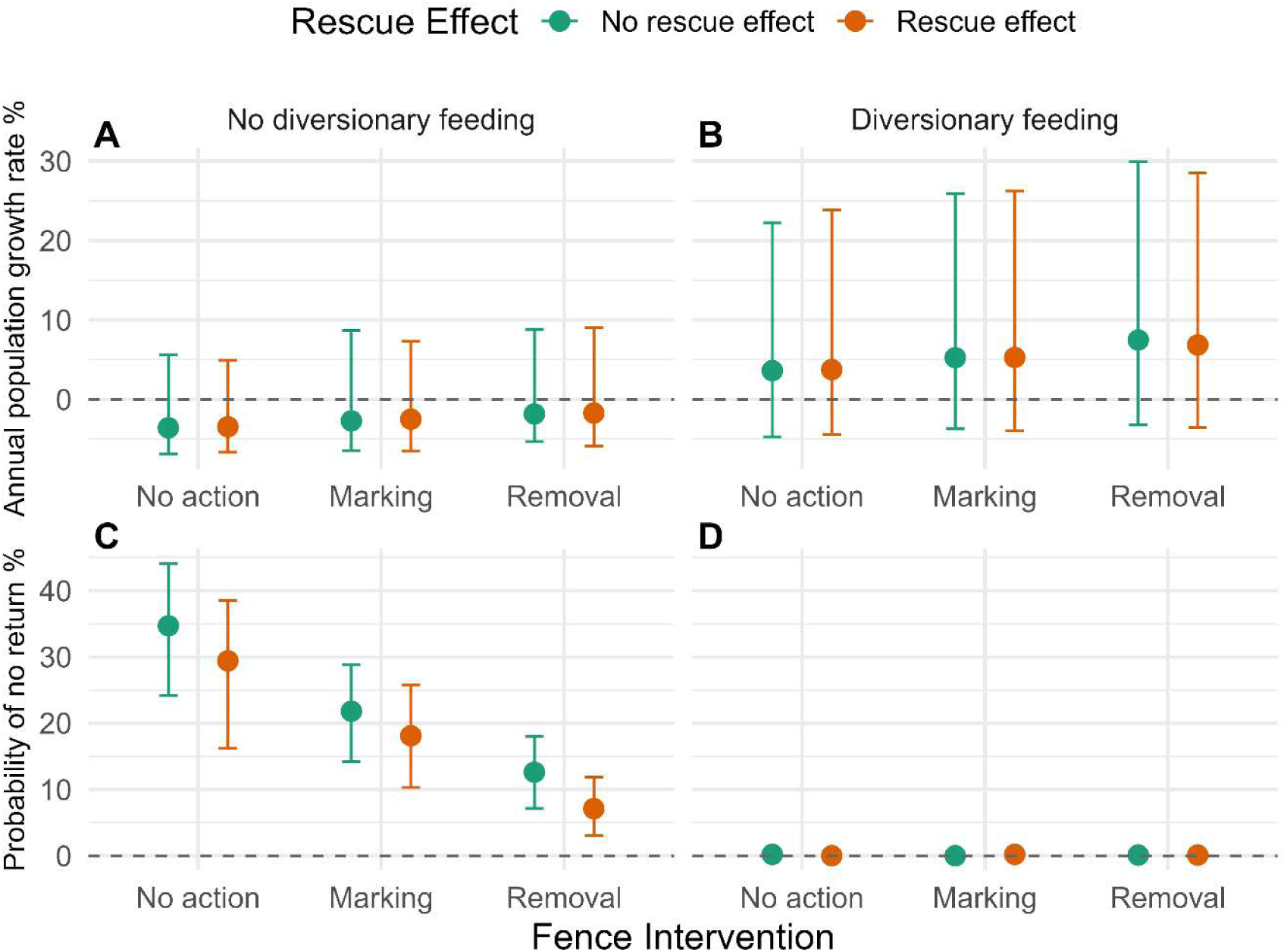
Lower bounds of management intervention effectiveness (diversionary feeding increasing reproductive rate; fence marking or removal improving survival) based on projections under scenarios with and without rescue effects. A, B: Median annual total population growth (2023-2033). C, D: Probability-of-no-return in 2060, defined as the probability of the population falling below 75 total female capercaillie.

## 4 Discussion

By integrating multiple data sources with biological hypotheses, our approach supports evidence-based conservation management under uncertainty for threatened populations such as the Scottish capercaillie. We improved the precision of population estimates, identified key environmental drivers, and evaluated proposed management actions. Our findings indicate that reversing declines may be achievable with available mitigation actions, although any recovery is likely to be gradual and would require sustained management and monitoring.

### 4.1 Population Monitoring

Integrating data sources markedly improved population estimate precision by up to 49.4% (Fig. 3) compared to standalone national surveys (Table 1). The estimated proportion of males also closely matched independent observations from national survey (Fig. 3B), adding confidence in the model’s demographic outputs.

Our status-quo simulation estimated extinction probabilities of 0% in 2045, and 1.6% in 2070, versus 23% and 95% in Baines & Aebischer (2023), likely reflecting their omission of observation error, which can bias extinction risk estimates (Baines & Aebischer, 2023). A key source of such bias is imperfect detection in lek counts, due to counting error and incomplete male attendance. We estimated only ≈30% males were observed at leks (Section 3.1; Table S2), substantially fewer than the commonly assumed 50% (Watson & Moss, 2008) and consistent with recent genetic-based assessments (Aleix-Mata et al., 2024; 0.34, 95% CI:0.26-0.43). This underscores the limitations of using lek counts as absolute indicators of population size (Baines & Aebischer, 2023; Metcalfe et al., 2022; Scottish Gamekeepers Association, 2024). Attendance likely varies with environmental and ecological variables such as weather and population density (Wann et al., 2019), warranting further empirical work to quantify these drivers and incorporate variability into models.

Improved precision can inform adaptive management by detecting population-level changes as new demographic data become available. This capability now supports evaluation of ongoing actions under the Capercaillie Emergency Plan (NatureScot & Cairngorms National Park Authority, 2024) and could inform similar conservation efforts for capercaillie populations across Europe (Coppes et al., 2016; Gil et al., 2020; Jiménez et al., 2022). More broadly, our results demonstrate the value of IPMs for threatened species conservation, particularly where monitoring data are biased or incomplete (e.g., lek counts). IPMs can refine population estimates, fill data gaps, and estimate latent parameters (e.g., detectability, proportion of males) that are challenging to measure directly (Saunders et al., 2019).

### 4.2 Drivers of population dynamics

Ground-nesting birds across Europe are experiencing declines (McMahon et al., 2024). For capercaillie in Scotland, we found strong evidence that fluctuations in voles (the preferred prey for capercaillie mesopredators) correlate with reproductive rate (Table 3). In years of low vole abundance, prey-switching increases nest predation (Kjellander & Nordström, 2003; Marcstrom et al., 1988; Wegge & Storaas, 1989). We hence provide strong evidence for the alternative prey hypothesis (Kjellander & Nordström, 2003; Marcstrom et al., 1988; Wegge & Storaas, 1989), consistent with patterns observed in Fennoscandia (Hilde et al., 2024). Therefore, dampened vole cycles across Europe over the past two decades, likely driven by changing winter conditions (Cornulier et al 2013), may have resulted in sustained high predation pressure on capercaillies in Scotland and beyond.

Pine martens, recovering in B&S since the 1990s (Summers, 2018), have been implicated in recent poor capercaillie breeding success (Baines et al., 2016), yet densities were stable between 2012-2022 (Hobson et al., 2025). Thus, any contribution of marten predation would not have worsened in recent decades. Badgers (*Meles meles*) likely expanded their range between 2006-2009 and 2022-2025 (Connolly et al., 2025), though they are five times less likely to predate nests than pine martens (Bamber et al., 2024). Artificial nest trials in 2024 showed overall predation rates ≈65%, consistent with those in Fennoscandia (Bamber et al., 2024).

Climate stressors may be one root cause of the capercaillie population vulnerability to predation (Wegge & Kastdalen, 2007). We found April warming and pre-breeding precipitation influenced reproductive rate (Table 3), consistent with previous studies (Baines et al., 2011; Coppes et al., 2021; Moss et al., 2001). Increased rainfall may impair foraging and reduce female body condition, leading to lower breeding success or chick viability (Watson & Moss, 2008), whereas a larger temperature rise in early April may stimulate plant growth, improving hen nutrition, body condition and consequently chick survival (Baines et al., 2011; Moss et al., 2001). Pre-breeding precipitation showed no temporal trend, suggesting conditions may already be suboptimal. In contrast to Moss et al., (2001), we detected a positive trend in April warming, which may have supported capercaillie dynamics in recent decades, though insufficient to offset other pressures such as predation. Unlike Moss et al. (2001) and Slagsvold & Grasaas (1979), we found no effect of rainfall during chick rearing (June rain days; Table 2) on reproductive rate (Table 3), agreeing with Wegge et al. (2022), possibly because other factors (e.g., predation pressure) now dominate variation.

Some drivers of decline (e.g., climate), may not be directly mitigable. In such cases, conservation efforts must focus on measures improving vital rates or indirectly influencing underlying drivers. For example, diversionary feeding may reduce nest predation by providing alternative food to prey-switching mesopredators offering a non-lethal strategy that avoids conflict with protected species such as pine martens (Bamber et al., 2024). Our model provided a tool that was able to evaluate these interventions, facilitating informed management decisions.

### 4.3 Management action effectiveness and recommendations

We evaluated three key management interventions: diversionary feeding, and fence removal or marking (Fig. 4; Fig. 5). Although fences were not identified as an important driver, this may reflect poor-quality fence data (see Section 4.4), which challenges the detection of interannual population impacts. Dedicated studies have demonstrated Capercaillie mortality due to fence collisions (Baines & Andrew, 2003; Baines & Summers, 1997; Moss, 2001). Our results indicate that no further deterioration in demographic rates can be endured by the population (Fig. 4), reinforcing the need for urgent action. We found that diversionary feeding alone was more effective than fence management and, if widely applied, may be sufficient to reverse declines, but improvements may highly uncertain and slow (Fig. 5B). Combining diversionary feeding and fence removal yielded the most favourable population outcomes overall (Fig. 5B).

Simulations assumed annual diversionary feeding across the entire B&S region. Although broadly supported, diversionary feeding has not yet replaced lethal predator control in all Scottish estates (Bamber et al., 2025a), which could limit its effectiveness. We also assumed uniform survival gains in space, though the effectiveness of fence interventions probably varies depending on the availability of fences to remove, fence type, marking type, visibility, and habitat (Baines & Andrew, 2003). Our estimates are based on a single marking type (orange netting) with wide confidence intervals (Baines & Andrew, 2003) and represent percentage increases from our baseline female survival rate (0.70), higher than the pre-mitigation rate reported by Moss et al. (2000; 0.63). Therefore, the benefit of further fence interventions is potentially overestimated.

These finding suggest that combining diversionary feeding and fence removal offers a promising route to recovery, but success will likely depend on spatiotemporal deployment, stakeholder engagement, and continued monitoring of their impact at the population-level.

### 4.4 Data limitations and model developments

Predator effects were particularly difficult to assess. Voles may act as a proxy for generalist predator pressure, but we were not able to directly test the influence of specific predators, as we lacked time series data for some species (e.g., badgers, foxes), and for pine martens had insufficient data for model convergence. Vole data were also temporally limited in B&S, so we reconstructed values based on assumed synchrony with Kielder vole cycles; supported by overlapping vole data, regional synchrony in vole dynamics (Cornulier et al., 2013), and broad correspondence with observed reproductive rate (Fig. S9). Pine marten scat counts are uncalibrated and may not reflect true abundance (Baines et al., 2013); calibrated approaches such as DNA-based spatially explicit capture-recapture (SECR) methods are preferable (Hobson et al., 2025). Even with better data, strong correlations among key covariates in boreal forest grouse systems may complicate causal inference (Wegge et al., 2022). More research is required to investigate confounding of field vole abundance with other habitat and environmental variables in Scotland.

Fence impacts were not detected in our analysis, likely due to data limitations. Our fence density covariate layer varied only spatially, because we lacked temporal information, although densities of fences declined from 1990 onwards. Assembling a comprehensive spatiotemporal dataset of fence removals and markings as well as newly erected fences would require broader data access and substantial archival data collation effort but could improve future assessments of management effectiveness and mortality attribution.

Monitoring practices have also evolved: brood counts with trained dogs are now largely discontinued due to disturbance concerns (Bamber et al., 2023; Metcalfe et al., 2022), while camera trap data have been collected to monitor breeding success since 2021 (Bamber et al., 2023). Non-invasive genetic sampling may also improve species detection rates (Augustine et al., 2020). The IPM framework allows integration of such new data sources, accounting for their distinct observation errors. We compared IPM reproductive rate estimates with those from camera trap data (Bamber et al., 2023, 2025b), finding closer agreement with chicks/adult hen than chicks/hen (Fig. S11; SI 8). We hypothesise, this may reflect lower dust bath use by 14-month hens or their use of unmonitored sites.

Our model did not attempt to estimate density dependence because lek sizes have probably remained below carrying capacity during the study period. However, estimating its strength will become important and feasible if the population begins to recover (Ginzburg et al., 1990).

Connectivity between subpopulations was also omitted because the population is declining and there was no evidence of asymmetric exchanges via dispersal; this also resulted in conservative extinction risk estimates by omitting the possibility of rescue effects at the stage of model fitting. Including rescue effects at the projection stage slightly reduced median long-term extinction risk, but credible bands overlapped with the non-rescue scenario and short-term growth rates were unaffected. Incorporating migration between leks could improve ecological realism in future models (Matthiopoulos et al., 2005), though implementation remains challenging due to convergence issues (e.g., when estimating female dispersal with sparse demographic data).

Beyond structural improvements, our framework could test scenarios such as population reinforcement (NatureScot & Cairngorms National Park Authority, 2024), continued dampened vole cycles, or projected climate impacts under alternative management strategies, and could be scaled up to the entire Scottish range. Future developments could directly quantify management impacts to support formal, iterative adaptive management (Nichols et al., 2007), identify minimum data requirements for reliable predictions by testing model performance under systematically reduced datasets (Keenan et al., 2013), and optimise spatial and temporal intervention deployment for efficient and cost-effective monitoring and management.

### 4.5 Conclusion

Our integrated approach provides a practical and flexible tool for identifying impacts of environmental drivers, monitoring population responses to management, and simulating interventions. Here, it is applied in support of the Capercaillie Emergency Plan (NatureScot & Cairngorms National Park Authority, 2024). More broadly, this approach illustrates how integrated modelling can inform evidence-based decision-making in complex conservation contexts, providing a flexible framework for intervention planning across species facing multifactorial declines.

## Acknowledgements

We thank the Royal Society for the Protection of Birds (RSPB), the Game and Wildlife Conservation Trust (GWCT), Forestry and Land Scotland (FLS), NatureScot, and Wildland for providing access to brood and lek count data. We thank University of Aberdeen and Forestry and Land Scotland for supplying vole data and the RSPB for access to fence data. We are grateful to Helen Gray (RSPB) for facilitating data access and mapping approximate capercaillie forest areas, and to Scott Newey (GWCT) for facilitating data access and providing helpful comments on an earlier manuscript draft. Xavier Lambin received a Leverhulme fellowship (RF-2024-363).

## Data Availability

Data and code supporting the findings of this study will be archived on Zenodo and made publicly available upon acceptance of the manuscript for publication in a peer-reviewed journal. A DOI will be provided at that time.

## Author Contributions

HIN: conceptualisation, formal analysis (lead), methodology (equal), visualisation, Writing – original draft preparation, writing – review & editing. JM: conceptualisation (initial), formal analysis (supporting), methodology (equal), supervision, writing - review & editing. FB: formal analysis (supporting), writing-review & editing, JAB: investigation, writing-review & editing, MD: conceptualisation (initial), investigation, resources. SE: conceptualisation (initial), writing – review and editing. KF: investigation, writing – review & editing. SH: conceptualisation (initial). KK: resources, writing-review & editing. XL: investigation, writing – review & editing, RM: conceptualisation, writing – review and editing. CS: writing – review & editing. LJZ: methodology (supporting), writing – review & editing.

