## Supplementary Information for "Applying integrated population models to guide conservation planning: Western Capercaillie in Scotland"

ψ Deceased

1. Testing for linear temporal trends in data, covariates and model estimates

For each datum, covariate, or model estimate, *x*, we fitted a linear model of the form $x \sim$ year using the lm function in R (R Core Team, 2023) . Model diagnostics included checks for:

- Linearity between predictors and response
- Homoscedasticity
- Normality of residuals
- Autocorrelation in residuals
- Influential observations

*Reproductive rate*

We tested for temporal trends in observed reproductive rate (from data), modelled reproductive rate (chicks per hen and chicks per adult hen) and the temporal random effect in reproductive rate. No significant temporal trends were detected (Table S1). Model for observed reproductive rate, estimated reproductive rate (chicks per adult hen) and temporal random effect (in reproductive rate) did not meet normality assumption, so we calculated robust Newey–West standard errors and p-value. Trends remained non-significant.

*Weather covariates*

We tested for temporal trends in each of the five climate covariates: pre-copulation precipitation, April temperature, snow cover, April warming index and June precipitation days.

No significant temporal trends were detected for any covariate (Table S1), apart from April warming index (see Section 2.1.2 for more details). All models met diagnostic assumptions, except for the snow cover model, which exhibited residual autocorrelation. To address this, we refitted the model using a linear mixed-effects approach with a first-order autocorrelation structure (corAR1) via the lme function (nlme; J. Pinheiro et al., 2024; J. C. Pinheiro & Bates, 2000) which resolved the autocorrelation issue. The trend remained non-significant.

### 2. Snow cover datum prediction

To estimate snow cover (Table 2; Section 2.1.2) for the year 2023, we fitted a Generalised Additive Model (GAM; S. N. Wood, 2017) of the form snow cover ~ s(year), (package *mgcv*; S. Wood, 2025). The smooth term for year allowed for non-linear temporal trajectories. The model was fitted using Restricted Maximum Likelihood (REML) to reduce the risk of overfitting and improve the reliability of predictions near the boundary of the observed data range. We then used the fitted model to estimate snow cover for 2023.

### 3. Survival between lek counts and national surveys

National surveys were done in winter, their midpoint (January) being about eight months after the lek counts in May. To estimate the number of individuals (adults and juveniles) surviving between lek counts and surveys, we applied the same model forms used in the process model for adult survival and juvenile recruitment over two thirds of a year.

Surviving adults were modelled as:

- Female adults: $F_{i,t+2/3} = B((1+2\varphi_{F\,i,t})/3, F_{i,t})$
- Male adults: $M_{i,t+2/3} = B((1+2\varphi_{M\,i,t})/3, M_{i,t})$

Surviving juveniles were modelled as:

- Female juveniles: $f_{0\,i,t+2/3} = Pois(C_{F\,i,t}\varphi_{F0a\,i,t+2/3}), f_{1\,i,t+2/3} = Pois(C_{F\,i,t-1}\varphi_{F1a\,i,t+2/3})$
- Male juveniles: $m_{0\,i,t+2/3} = Pois(C_{M\,i,t}\varphi_{M0a\,i,t+2/3}), m_{1\,i,t+2/3} = Pois(C_{M\,i,t-1}\varphi_{M1a\,i,t+2/3}), m_{2\,i,t+2/3} = Pois(C_{M\,i,t-2}\varphi_{M2a\,i,t+2/3})$

Where the cumulative survival terms over two-thirds of the year were defined as:

- $\varphi_{F0a\,i,t+2/3} = \frac{(1+2\varphi_{jF\,i,t})}{3}$
- $\varphi_{F1a\,i,t+2/3} = \frac{(1+2\varphi_{F\,i,t})}{3}\varphi_{jF\,i,t-1}$
- $\varphi_{M0a\,i,t+2/3} = \frac{(1+2\varphi_{jM\,i,t})}{3}$
- $\varphi_{M1a\,i,t+2/3} = \frac{(1+2\varphi_{M\,i,t})}{3}\varphi_{jM\,i,t-1}$
- $\varphi_{M2a\,i,t+2/3} = \frac{(1+2\varphi_{M\,i,t})}{3}\varphi_{M\,i,t-1}\varphi_{jM\,i,t-2}$

4. Priors

To initialise the model in the first year (1990), we used gamma-distributed priors for population size, based on the mean and standard deviation of 1989 lek counts (Picozzi et al., 1992). Gamma prior for the baseline reproductive rate ($b_0$) was informed by Fletcher & Baines, 2020, with a standard deviation of 1 to allow moderate uncertainty. Survivals ($\theta_x$) were informed by Moss et al., 2000, with standard deviations derived from confidence intervals. First-year survival, $\theta_{jF}$, was parametrised as a proportion of adult survival $\theta_F$, to constrain the parameter space and improve model convergence , such that $\text{logit}^{-1}(\theta_{jF}) = r\text{logit}^{-1}(\theta_F)$, where r is the ratio between estimated median juvenile and female survival values (0.5÷0.63) from Moss et al 2000. The prior for the proportion of male chicks ($p_C$) was specified as a beta distribution with mean 0.41 and standard deviation 0.05, reflecting evidence of female-biased broods, likely due to lower male chick survival when reproductive rate is low (Moss et al., 2000).

The proportion of the total male population observed at leks ($p_l$) was modelled using a beta prior centred at 0.5, based on estimates from (Watson & Moss, 2008) with a standard deviation of 0.1, to allow a plausible detection range from approximately 0.2 to 0.8. National survey data is scaled to account for imperfect detectability (Ewing et al., 2012; Wilkinson et al., 2018, 2023) therefore, we assumed surveys approximately represent the total population scaled by detection probability $p_s$. We centred $p_s$ on 1, with a standard deviation of approximately 0.05 allowing for deviations between approximately 0.8 and 1.2 to reflect uncertainty.

We applied stronger constraints on random effect variances in survival than in reproduction, both to aid model convergence and because survival is typically a more buffered vital rate (Hilde et al., 2020). Random effects were modelled using gamma priors. For reproductive rate, the random effect was constrained to allow an increase in reproductive rate up to three times the baseline rate. For survival, random effect priors allowed for deviations of up to 2% from the baseline rate.

Priors for covariate coefficients were set as uninformative in magnitude, and the effect direction (+ve or -ve) was informed by the literature (Table 2). For example, the effect of fences on first-year survival was *a priori* assumed to be negative. Snow cover was included as a proxy for unavailable snow melt data (Table 2). Therefore, we were unsure of the expected direction of its effect, so snow cover was assigned a normal prior centred on zero, allowing for both positive and negative influence.

The standard deviation representing expected variability in Badenoch and Strathspey (B&S) vole indices around the Kielder vole index ($\sigma_V$) was given a Gamma prior (shape = 1, rate = 50), which strongly constrains deviations from the Kielder index, corresponding to an expected deviation up to ≈5% relative to the mean Kielder unstandardised index. This reflects the limited evidence for large-scale divergence between regions and ensures that the vole covariate does not absorb unexplained variation in reproductive rate unrelated to vole dynamics.

See Table S2 for full prior specifications.

### 5. Modelling Pine Martens

Pine marten data were very sparse, with index data available for only three years (1995, 2009, 2013), across a maximum of 7 forests (Section 2.1.2). Although there is evidence of a historical increase in pine marten population size (Baines et al., 2016) (see main manuscript section 4.4 on data limitations), data from spatially explicit capture-recapture using DNA-derived individual identification indicate a plateau in density between 2012 and 2022 (Hobson et al., 2025).

Given these patterns, we modelled pine martens using a density-dependent growth process, where $a_0$ is the growth rate parameter, and $a_1$ the density-dependent parameter:

$$P_{m\,rate\,f,t} = P_{m\,f,t}\exp\left(a_0 - a_1 P_{m\,f,t}\right)$$

The observation process was modelled as a truncated normal distribution (lower bound=0), with mean equal to the predicted pine marten rate and standard deviation, $\sigma_{PM}$.

$$P_{m\ \ f,t+1} \sim Norm(P_{m\,rate\,f,t}, \sigma_{PM})$$

The priors were chosen to constrain the reconstructed pine marten indices due to limited data availability to inform them (Table S2).

Due to limited data availability, models including pine martens as covariates failed to converge. The spatial structure in our model allows some level of space-for-time substitution (Pickett, 1989) to capture the influence of covariates with short time series, but in this case the data were too few. As a result, pine martens were excluded from model selection.

### 6. State-space model with covariate selection

We assessed covariate support using indicator variable selection, also known as a spike and slab approach (O'Hara & Sillanpää, 2009), following the Kuo and Mallick method (Kuo & Mallick, 1998). This approach incorporates uncertainty in variable inclusion directly into the model, allowing the data to inform which covariates are supported, so avoiding exhaustive model comparison (O'Hara & Sillanpää, 2009). For each covariate, $z$, we specified an indicator variable, $I_z \sim \text{Bernoulli}(p_I)$, with $p_I \sim \text{Beta}(2,4)$ (mean≈0.3; Table S2), a weakly informative prior reflecting that several covariates may be influential. Covariate support was assessed using posterior inclusion probabilities (PIPs), defined as the posterior mean of the indicator variable $I_z$, and summarized influential covariates using the median probability model (MPM; (Barbieri & Berger, 2004), which includes covariates with PIP $> 0.5$.

The model was implemented in JAGS (Plummer, 2003) and interfaced with R version 4.3.1 (R Core Team, 2023) using the *runjags* package version 2.2.2-4 (Denwood, 2016). We ran 4 chains with a burn-in of 1,660,000, sample of 10000 and thinning interval of 10. Convergence was assessed using the Gelman-Rubin diagnostic (Gelman & Rubin, 1992) and visual inspection of trace plots.

7. Calculating percentage effectiveness of management actions

We estimated the percentage increase in survival due to fence removal or marking from survival estimates (Moss et al., 2000) and in breeding success due to diversionary feeding using a bootstrapping approach with 10,000 samples. All standard deviations were calculated from 95% confidence intervals (CIs), assuming a normal distribution.

*Diversionary Feeding*

We sampled from normal distributions defined by the median and estimated standard deviation of breeding success in August (Bamber et al., 2025), with ($b_I$) and without ($b_C$) diversionary feeding:

$$Increase\ b = \frac{b_I - b_C}{b_I} \times 100$$

From the resulting distribution of percentage increases, we extracted the median and 95% confidence intervals.

*Complete Fence Removal*

We sampled from normal distributions representing:

- Joint adult survival including fence mortality (median=0.72) (Moss et al., 2000)
- Increase in survival in the absence of fence mortality (median=0.08) (Moss et al., 2000)

The percentage increase was calculated as:

$$\text{Increase } s = \frac{s_r}{s_0} \times 100$$

As above, we extracted the median and 95% confidence intervals from the bootstrapped distribution.

*All Fences Marked*

We sampled from a normal distribution using the reported median and estimated standard deviation of percentage collision reduction due to fence marking (Baines & Andrew, 2003). This multiplied the bootstrapped survival increase from complete fence removal distribution, and the resulting distribution was used to extract the median and 95% confidence intervals.

8. Comparison of reproductive rate estimates derived from the model and camera trap data

This was an exploratory comparison to investigate potential similarities and differences between modelled reproductive rate and camera trap-derived reproductive rate (Fig. S11), which could inform future integration of this monitoring approach into the IPM. Differences may reflect age-class variation in dust bath use or sampling bias due to trap placement at active sites, though no studies have investigated this. Dust bathing is primarily used in birds for parasite control and feather maintenance (Bamber et al., 2023; Olsson & Keeling, 2005).

Figure and Table Legends:

Figure S1: Number of leks where male capercaillie were counted per year 1990-2020.

Figure S2: Number of forests per year in which brood counts were conducted 1990-2020 (see Section 2.1.1; Figure 1).

Figure S3: Time series of standardised pre-copulation precipitation from 1990 to 2020 (see Table 2 for definition).

Figure S4: Time series of standardised April warming index from 1990 to 2020 (Table 2 for definition).

Figure S5: Time series of standardised June precipitation days from 1990 to 2020 (Table 2 for definition).

Figure S6: Time series of standardised spring snow cover from 1990 to 2020 (Table 2 for definition).

Figure S7: Time series of standardised April temperature from 1990 to 2020 (Table 2 for definition).

Figure S8: Standardised Badenoch and Strathspey (B&S) and Kielder vole data plotted together for visual comparison.

Figure S9. Standardised Kielder vole indices and standardised observed capercaillie reproductive rate (chicks per hen) plotted together for visual comparison.

Figure S10. Standardised Kielder vole indices and standardised capercaillie reproductive rate (chicks per hen), as observed and as modelled respectively, plotted together for visual comparison.

Figure S11. Reproductive rate as estimated by camera traps vs model estimates.

Table S1: Results from linear models testing for temporal trends in observed reproductive rate, modelled reproductive rate (chicks per hen and chicks per adult hen), the temporal random effect in reproductive rate and weather covariates. Values shown are median trend estimates, 95% confidence intervals (CIs) and p-values.

Table S2: Model symbols, parameters and variables including, where relevant, prior specifications and posterior summaries for the Bayesian state-space model with covariate selection. The table includes model symbols, descriptions, prior distributions, prior means and standard deviations (SDs), and posterior medians with upper and lower 95% credible intervals limits (CILs). Posterior estimates are not reported for parameters relating to pine martens, as these models did not converge. Posterior estimates for initial population size were not monitored, as this parameter was not central to inference and excluding it reduced model output size.

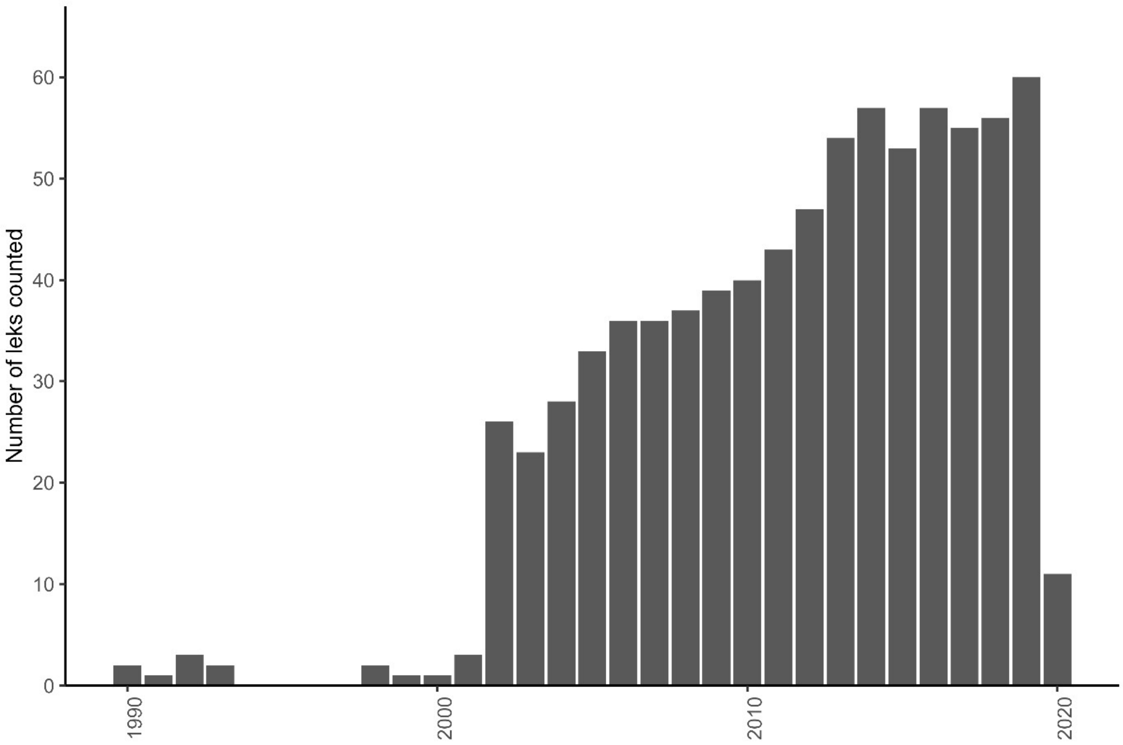


Figure S1

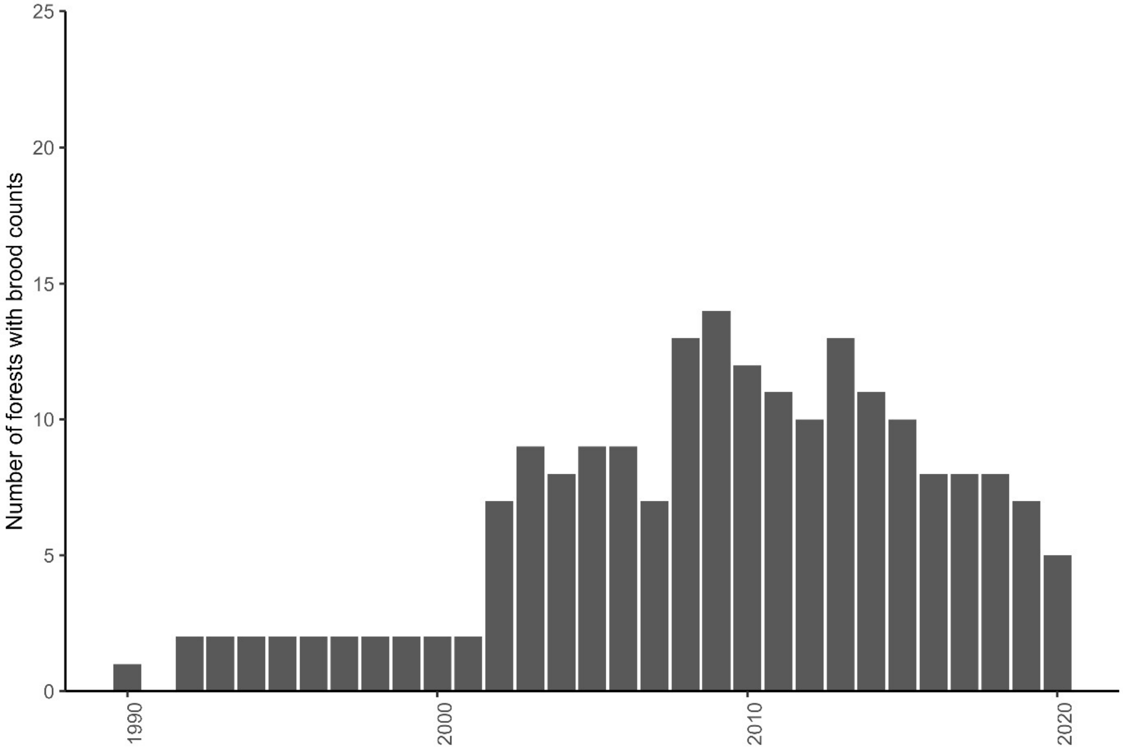


Figure S2

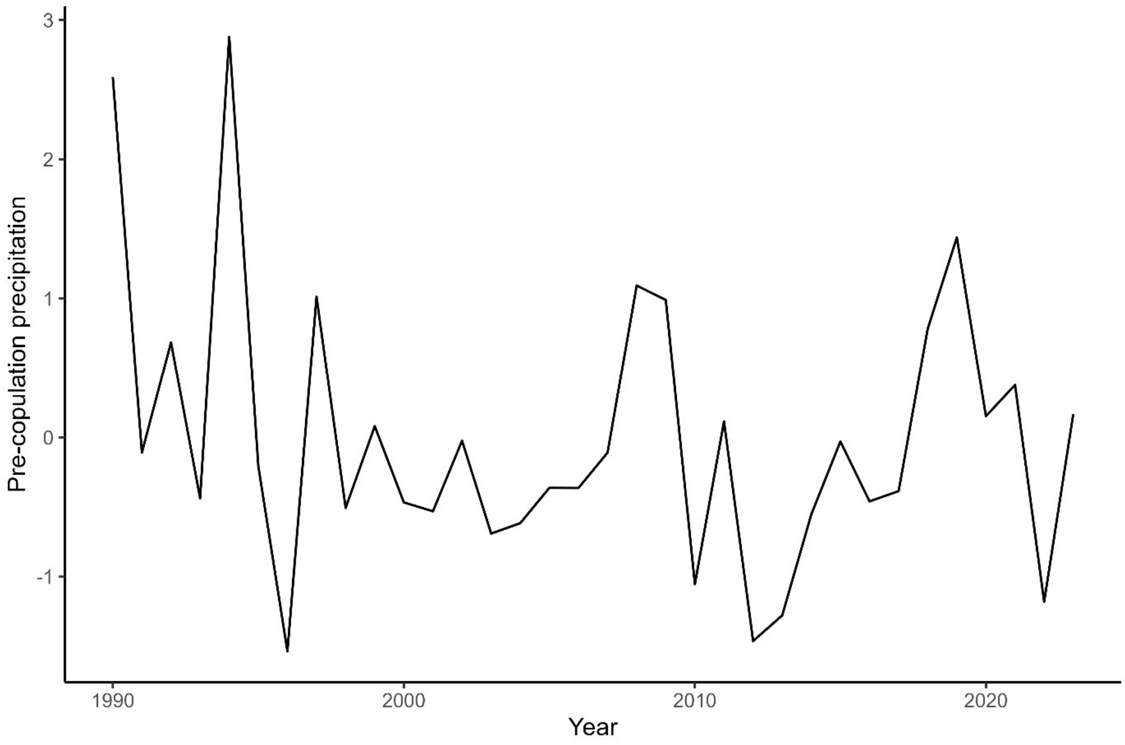


Figure S3

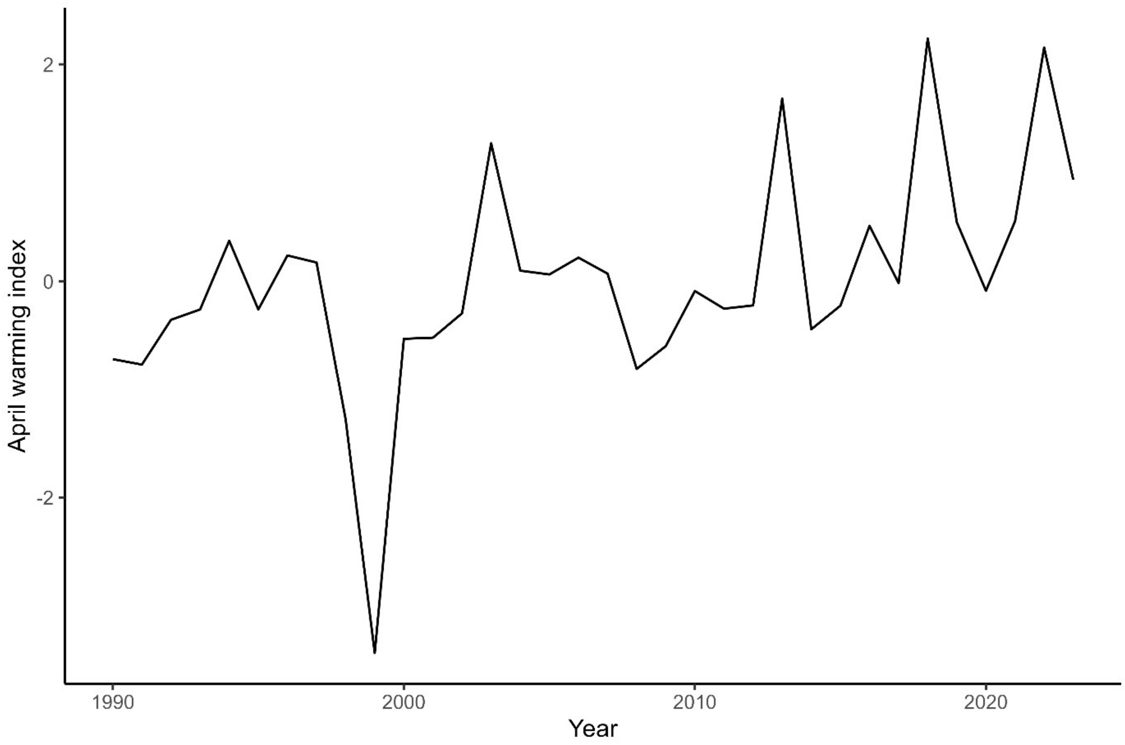


Figure S4

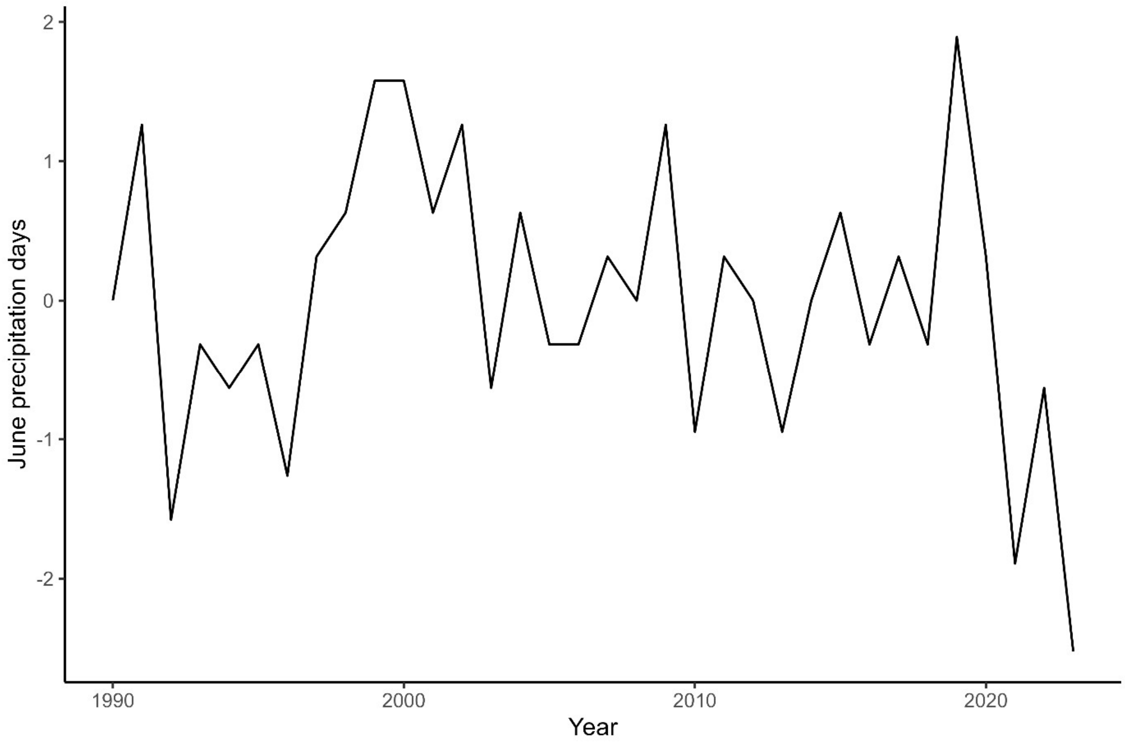


Figure S5

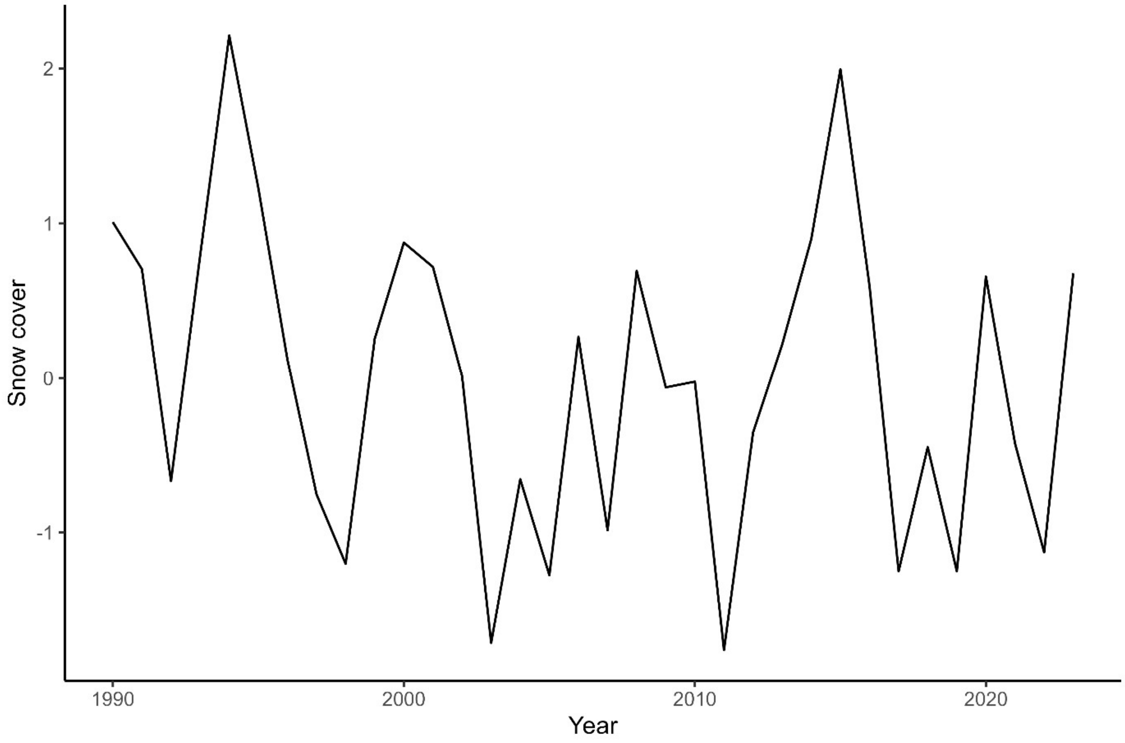


Figure S6

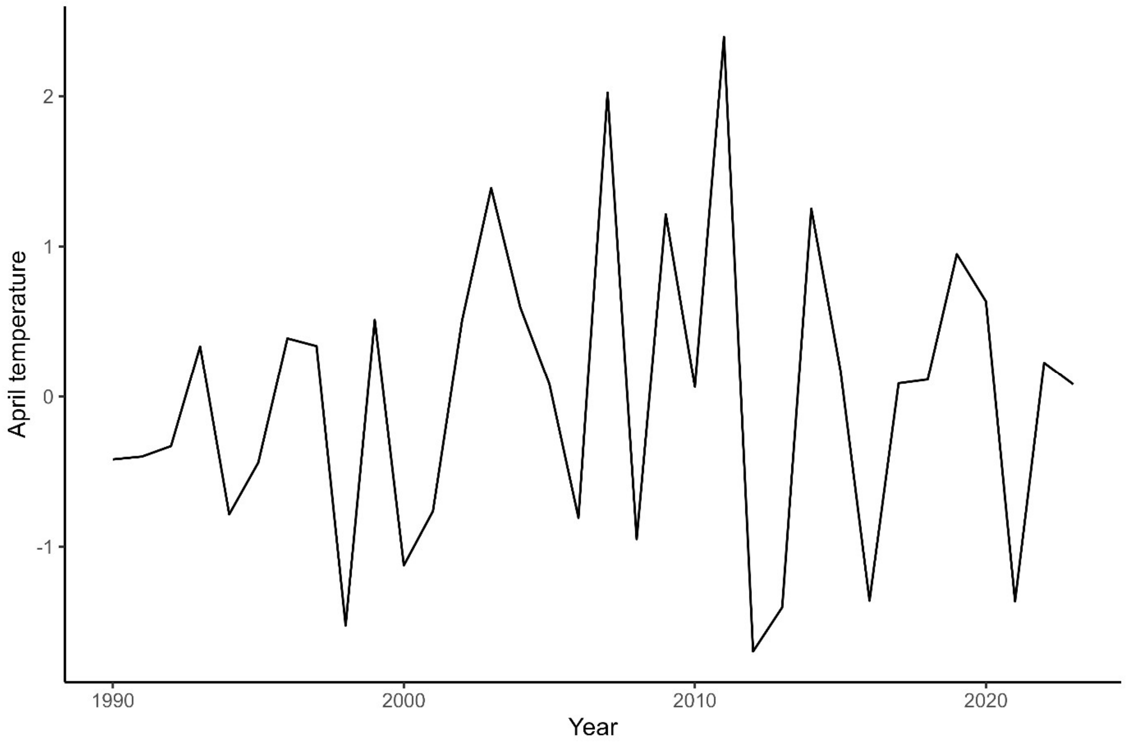


Figure S7

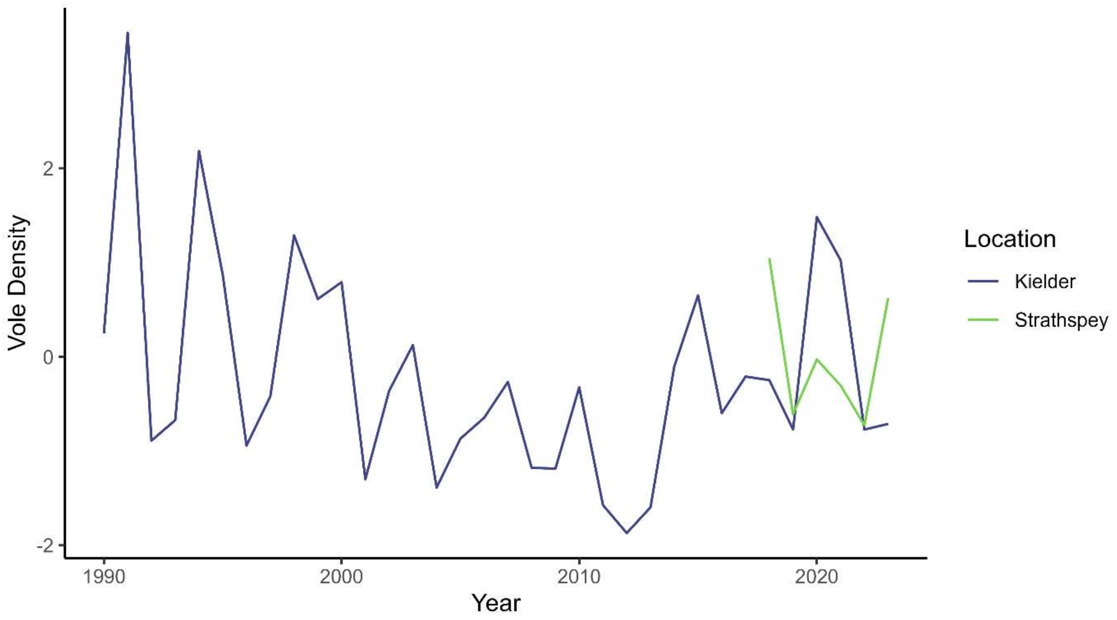


Figure S8

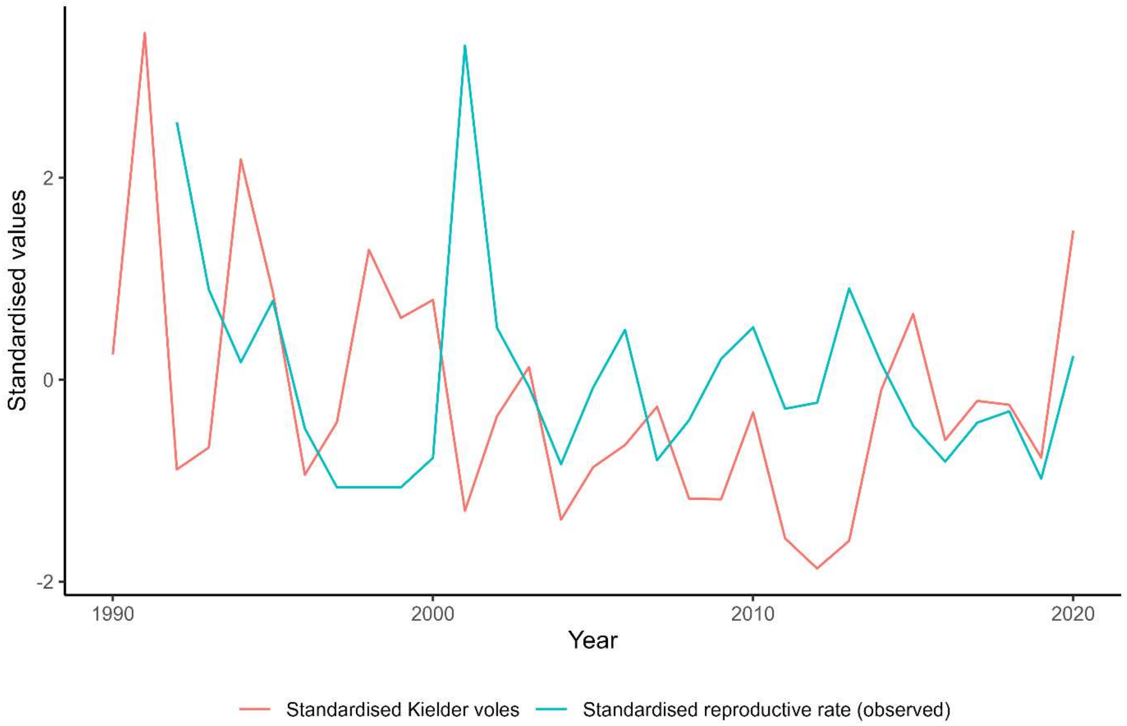


Figure S9

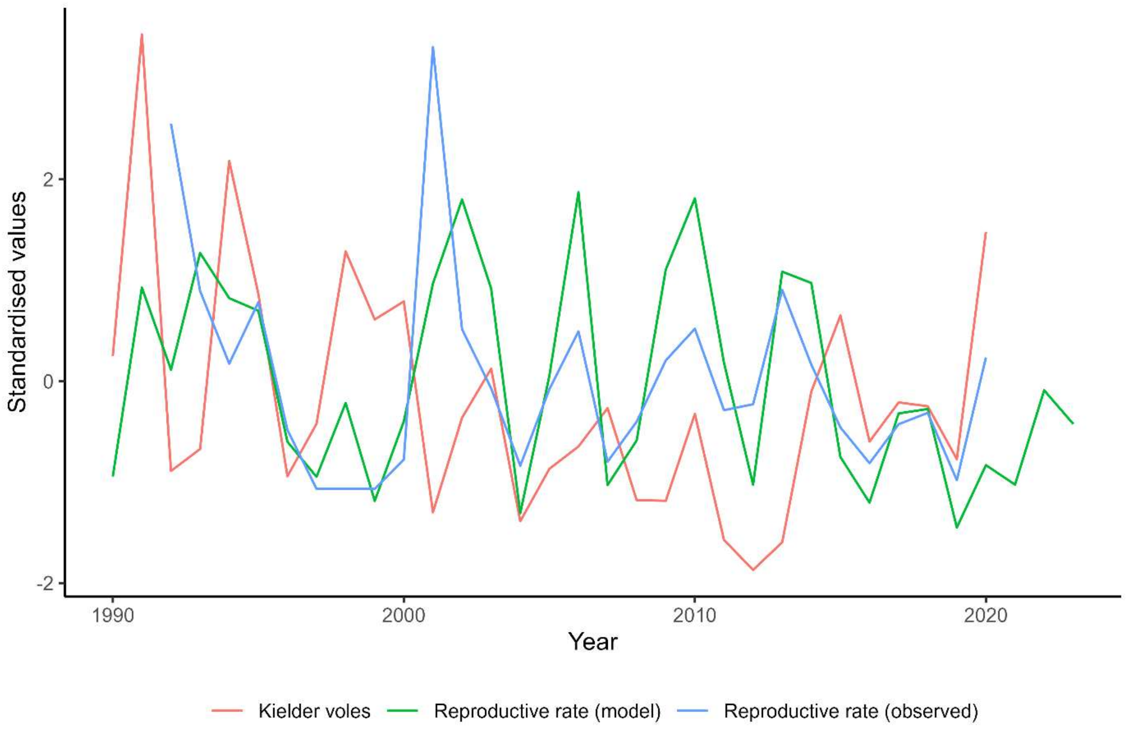


Figure S10

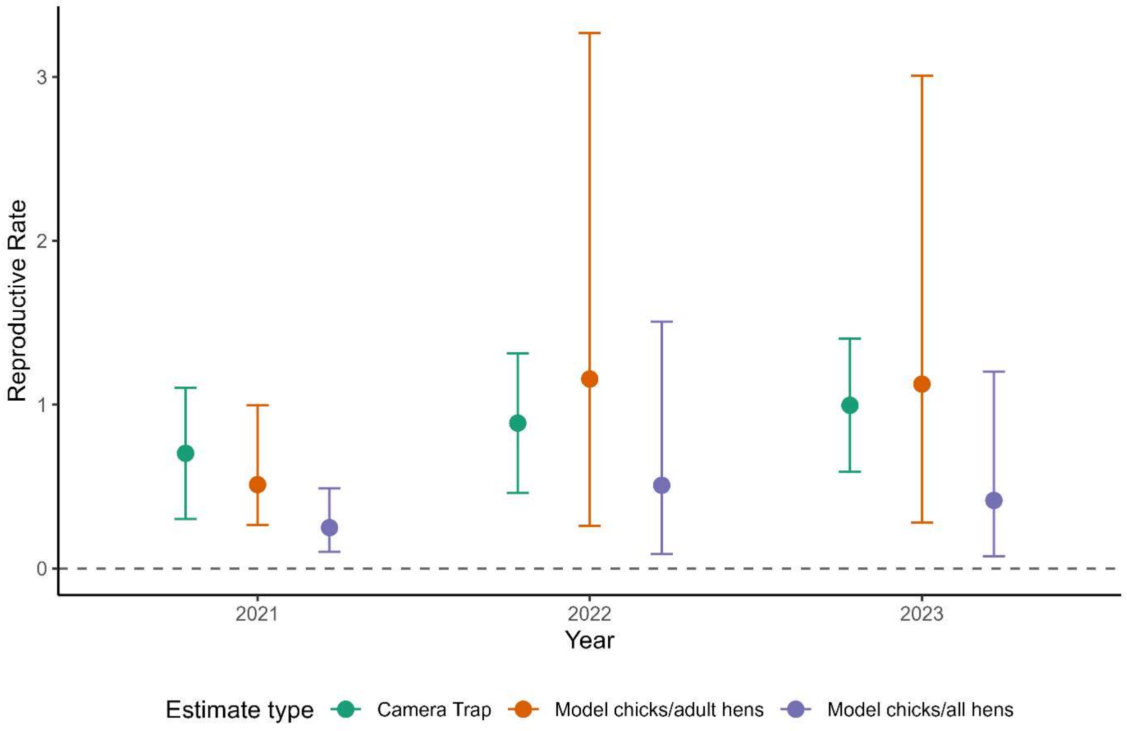

Reproductive Rate
3
2
1
0
2021
2022
2023
Year
Estimate type
Camera Trap
Model chicks/adult hens
Model chicks/all hens


Figure S11

| Covariate/data/model estimate | Median | Lower CI | Upper CI | p value |
|---|---|---|---|---|
| Observed reproductive rate | -0.0314 | -0.0766 | 0.0138 | 0.165 |
| Estimated reproductive rate (chicks/hen) | -0.0247 | -0.0599 | 0.0101 | 0.158 |
| Estimated reproductive rate (chicks/adult hen) | 0.00763 | -0.0284 | 0.0437 | 0.669 |
| Reproductive rate temporal random effect | 0.00593 | -0.0302 | 0.0420 | 0.740 |
| Pre-copulation precipitation | -0.0205 | -0.0559 | 0.0149 | 0.247 |
| April warming index | 0.0487 | 0.0170 | 0.0803 | 0.00367 |
| June precipitation days | -0.0175 | -0.0538 | 0.0174 | 0.305 |
| Snow cover | -0.0213 | -0.0698 | 0.0273 | 0.379 |
| April temperature | 0.0104 | -0.0255 | 0.0464 | 0.559 |

Table S1

| Symbol/Parameter/Variable | Description | Prior Distribution | Prior Mean | Prior SD | Posterior median | Lower CIL | Upper CIL |
|---|---|---|---|---|---|---|---|
| $i$ | Lek index $\in \{1, \dots, 67\}$ | NA | NA | NA | NA | NA | NA |
| $t$ | Year index $\in \{1, \dots, 34\}$ | NA | NA | NA | NA | NA | NA |
| $z$ | Covariate index in reproductive rate $\in \{1, \dots, 6\}$ | NA | NA | NA | NA | NA | NA |
| $x$ | Sex $\in \{F, M\}$ | NA | NA | NA | NA | NA | NA |
| $f$ | Forest index $\in \{1, \dots, 25\}$ | NA | NA | NA | NA | NA | NA |
| $\theta_F$ | Baseline female survival | Normal | 0.51 | 0.38 | 0.93 | 0.79 | 1.08 |
| $\theta_M$ | Baseline male survival | Normal | 1.47 | 0.5 | 1.73 | 1.46 | 1.98 |

| | | | | | | | |
|---|---|---|---|---|---|---|---|
| $\theta_j$ | Baseline juvenile survival written in terms of $\theta_F$ (SI 4) | NA | NA | NA | 0.27 | 0.17 | 0.36 |
| $b_0$ | Baseline reproductive rate | Normal | -1.39 | 1 | -0.28 | -0.47 | -0.10 |
| $p_C$ | Proportion of male chicks | Beta | 0.41 | 0.05 | 0.31 | 0.22 | 0.40 |
| $p_S$ | National survey detection probability | Gamma | 1.0 | 0.05 | 0.96 | 0.88 | 1.04 |
| $p_l$ | Proportion of males seen at leks | Beta | 0.5 | 0.1 | 0.34 | 0.28 | 0.40 |
| $p_I$ | Inclusion probability | Beta | 0.33 | 0.18 | 0.35 | 0.10 | 0.62 |
| $\sigma_V$ | Standard deviation of B&S voles relative to Kielder voles | Gamma | 0.02 | 0.02 | 1.08 | 0.94 | 1.26 |

| | | | | | | | |
|---|---|---|---|---|---|---|---|
| $a_0$ | Pine marten population growth rate | Gamma | 0.01 | 0.01 | | | |
| $a_1$ | Pine marten density-dependence | Gamma | 0.01 | 0.01 | | | |
| $\sigma_{PM}$ | Pine marten standard deviation | Gamma | 0.94 | 3.2 | | | |
| $\tau_{f\ i,1}$<br>$\tau_{m\ i,1}$ | Initial population prior | Gamma | 5.4 | 3.0 | | | |
| $\beta_1$ | Voles coefficient | +Gamma | 1 | 1 | Main manuscript Table 3 | Table 3 | Table 3 |
| $\beta_2$ | Pre-copulation precipitation coefficient | -Gamma | 1 | 1 | Table 3 | Table 3 | Table 3 |
| $\beta_3$ | April warming index coefficient | +Gamma | 1 | 1 | Table 3 | Table 3 | Table 3 |

| | | | | | | | |
|---|---|---|---|---|---|---|---|
| $\beta_4$ | June precipitation days coefficient | -Gamma | 1 | 1 | Table 3 | Table 3 | Table 3 |
| $\beta_5$ | Snow cover coefficient | Normal | 0 | 1 | Table 3 | Table 3 | Table 3 |
| $\beta_6$ | April temperature coefficient | -Gamma | 1 | 1 | Table 3 | Table 3 | Table 3 |
| $\beta_\gamma, \beta_{jF\gamma}, \beta_{jM\gamma}$ | Fence coefficients (adults, juvenile females and juvenile males) | -Gamma | 1 | 1 | Table 3 | Table 3 | Table 3 |
| $P_{tot\ i,t}$ | Total population size | NA | NA | NA | NA | NA | NA |
| $F$ | Adult females | NA | NA | NA | NA | NA | NA |
| $M$ | Adult males | NA | NA | NA | NA | NA | NA |
| $f_0, f_1$ | Subadult females | NA | NA | NA | NA | NA | NA |
| $m_0, m_1, m_2$ | Subadult males | NA | NA | NA | NA | NA | NA |
| $C_{F\ i,t}, C_{M\ i,t}$ | Female and male chicks | NA | NA | NA | NA | NA | NA |

| | | | | | | | |
|---|---|---|---|---|---|---|---|
| $S_{F\ i,t}, S_{M\ i,t}$ | Surviving adults | NA | NA | NA | NA | NA | NA |
| $\varphi_{F\ i,t}, \varphi_{M\ i,t}, \varphi_{jF\ i,t}, \varphi_{jM\ i,t}$ | Adult and juvenile survival probabilities | NA | NA | NA | NA | NA | NA |
| $\gamma_i$ | Fence density | NA | NA | NA | NA | NA | NA |
| $\delta_i, \delta_t$ | Lek and year random effects (survival) | NA | NA | NA | NA | NA | NA |
| $\mu_{i,t}$ | Reproductive rate (chicks/adult hen) | NA | NA | NA | NA | NA | NA |
| $\alpha_t, \alpha_i$ | Lek and year random effects (reproductive rate) | NA | NA | NA | NA | NA | NA |
| $P_{s\ t}$ | Observed national survey count | NA | NA | NA | NA | NA | NA |
| $\upsilon_t$ | Observed national survey precision | NA | NA | NA | NA | NA | NA |

| | | | | | | | |
|---|---|---|---|---|---|---|---|
| $l_{i,t}$ | Observed lek count | NA | NA | NA | NA | NA | NA |
| $p_{h\ f,t}$ | Proportion of hens per forest | NA | NA | NA | NA | NA | NA |
| $\mu_{f,t}$ | Forest area reproductive rate | NA | NA | NA | NA | NA | NA |
| $c_{f,t}$ | Observed chick counts | NA | NA | NA | NA | NA | NA |
| $h_{f,t}$ | Observed hen counts | NA | NA | NA | NA | NA | NA |
| $V_{f,t}$ | Standardised Badenoch and strathspey vole indices | NA | NA | NA | NA | NA | NA |
| $V_{K\ t}$ | Standardised Kielder vole index | NA | NA | NA | NA | NA | NA |
| $P_{m\ rate\ f,t}$ | | NA | NA | NA | NA | NA | NA |
| $P_{m\ f,t}$ | Pine marten scats | NA | NA | NA | NA | NA | NA |
| $I_z$ | Indicator variable | NA | NA | NA | NA | NA | NA |

Table S2
